# Live imaging of heart tube development in mouse reveals alternating phases of cardiac differentiation and morphogenesis

**DOI:** 10.1101/170522

**Authors:** Kenzo Ivanovitch, Susana Temiño, Miguel Torres

**Affiliations:** Developmental Biology Program, Centro Nacional de Investigaciones Cardiovasculares (CNIC), Madrid, Spain

**Author notes:** Current Address, The Francis Crick Institute, London, UK.

## Abstract

During vertebrate heart development two progenitor populations, first and second heart fields (FHF, SHF), sequentially contribute to longitudinal subdivisions of the heart tube (HT), with the FHF contributing the left ventricle and part of the atria, and the SHF the rest of the heart. Here we study the dynamics of cardiac differentiation and morphogenesis by tracking individual cells in live analysis of mouse embryos. We report that during an initial phase, FHF precursors differentiate rapidly to form a cardiac crescent, while limited morphogenesis takes place. In a second phase, no differentiation occurs while extensive morphogenesis, including splanchnic mesoderm sliding over the endoderm, results in HT formation. In a third phase, cardiac precursor differentiation resumes and contributes to SHF-derived regions and the dorsal closure of the HT. These results reveal tissue-level coordination between morphogenesis and differentiation during HT formation and provide a new framework to understand heart development.

## INTRODUCTION

The heart is the first organ to form and function during embryonic development. At embryonic day (E) 7.5, cardiac precursors in the splanchnic mesoderm (mesoderm apposed to the endoderm) differentiate into cardiomyocytes by assembling the contractile sarcomere machinery (Tyser et al., 2016) and form a bilateral structure known as the cardiac crescent (cc) in the mouse. Concomitant with foregut invagination, the cc swings inwards to become placed underneath the developing head folds. By a complex morphogenetic process, the cc subsequently transforms into an early heart tube (HT) initially opened dorsally, which by E8.25 has transformed into a closed and beating linear HT, also known as the primitive HT (Evans et al., 2010; Kelly et al., 2014).

The cc and early HT mainly give rise to the left ventricle (Zaffran et al., 2004). The right ventricle (RV), the outflow tract and most of the atria derive instead from cardiac progenitors located dorso-medially to the cc in the splanchnic mesoderm, that are progressively recruited at the poles of the HT at subsequent developmental stages (Cai et al., 2003; Kelly et al., 2001; Mjaatvedt et al., 2001; Waldo et al., 2001; Galli et al., 2008 135:1157-67). These findings led to the concept that cardiac mesodermal progenitors contain two populations of cells: the first heart field (FHF) precursors, recruited early in development to form the initial HT and mostly containing the LV primordium, and the second heart field (SHF), recruited later and elongating the HT (Buckingham et al., 2005).

Clonal analysis (Devine et al., 2014; Lescroart et al., 2014; Meilhac et al., 2004a) supports the idea that FHF and SHF precursors are two independent developmental fields with dedicated molecular pathways. Clonal analysis also showed that the SHF shares a common origin with the skeletal muscles of the head and neck within the pharyngeal mesoderm (Lescroart et al. 2010, Lescroart et al. 2015), further supporting differences between FHF and SHF populations. However, the existence of a common precursor between FHF and SHF was also reported in the early mouse embryo (Meilhac et al., 2004a) and other views suggest that the heart would form by a continuous differentiation process from a single population of cardiac precursors and only timing of recruitment would distinguish cells of the FHF and SHF (Abu-Issa et al., 2004; Moorman et al., 2007). In support of the latter, the typical marker of the SHF, Islet1 (Cai et al., 2003), is also transiently expressed in FHF precursors and must therefore be considered as widespread cardiac progenitor marker instead (Cai et al., 2003; Prall et al., 2007; Yuan and Schoenwolf, 2000). Whether the recruitment of cardiomyocytes from progenitors is a continuous process and how this coordinates with morphogenesis, however, has not been directly studied. This is partly because the spatial arrangement of progenitors and differentiated cardiomyocytes has so far been analyzed on fixed embryos (Cai et al., 2003; Spater et al., 2013) and the expression dynamics of genes reporting differentiation together with cell movements during HT morphogenesis have not been captured so far (Abu-Issa, 2014).

Here, we report the live-imaging and 3D+t cell tracking of HT formation in whole mouse embryos. Using this method, in conjunction with an Nkx2.5eGFP reporter line, which provides high level of GFP in differentiated cardiomyocytes, we studied the dynamics of cardiac field differentiation. During an initial phase, FHF cardiac precursors differentiate rapidly to form a cardiac crescent, while limited morphogenesis takes place. During a second phase, no differentiation events are detected and extensive morphogenesis, including splanchnic mesoderm sliding over the endoderm, results in HT formation. Finally, using an Isl1-Cre lineage tracing assay combined with live-imaging, we show that during a third phase, cardiac precursor differentiation resumes and contributes not only to the known SHF-derived regions but also to the dorsal aspect of the HT. These results show essential properties of FHF and SHF contribution to heart development and reveal tissue-level coordination between alternating phases of differentiation and morphogenesis during HT formation.

## RESULTS

### 3D static analysis of mouse HT formation

To assess how the initial cardiogenic region transforms into a HT and differentiates, we first analyzed *Nkx2.5^cre/+^*; *Rosa26Rtdtomato^+/−^* embryos, in which both cardiac precursors and cardiomyocytes are labeled (Stanley et al., 2002). Before cc differentiation, at early head fold stage (EHF, ~E7.5), the cardiogenic region is visualized as a flat horse shoe-shaped tdtomato+ mesodermal layer at the rostral border of the embryo (Figure 1A, A’, and Video 1). Figure 1-figure supplement-1 shows the criteria for embryo staging (Downs and Davies, 1993; Lawson, 2016). In the *Nkx2.5^cre/+^; Rosa26Rtdtomato^+/−^* embryos, tdtomato labeling is also observed in the endocardium and endothelial cells (Stanley et al., 2002) but not in the endoderm (Figure 1-figure supplement 2A, A’). We next studied the distribution of Cardiac troponin T (cTnnT), one of the first evident sarcomeric proteins to appear in the cardiac crescent (Tyser et al., 2016). At EHF stage (Figure 1B), while most embryos are negative for cTnnT expression, some embryos show weak cTnnT localization in subsets of cells (Figure 1-figure supplement 3A, A’). At a subsequent embryonic stage (~E7.7), cTnnT signal reveals the cc, which is folding inwards. During folding, the cTnnT signal increases. cTnnT+ cells are initially columnar epithelial cells and show apical localization of the tight junction component zona-occluden-1 (ZO-1) (Figure 1-figure supplement 3B, B’). During differentiation, cardiac precursors switch to a rounded shape (Linask et al., 1997) (Figure 1C, D) and separate from the endoderm, while maintaining a basal lamina at the endocardial side (inset in Figure 1D and Figure 2D). Morphogenetic changes starting at ~E8 subsequently lead to the formation of a hemi-tube whose major axis is transversal to the embryo A-P axis. We will refer to this stage as transversal HT (Figure 1E). Later, the tube adopts a more spherical shape, very similar to the linear HT but still open dorsally. We will refer to this stage as open HT (Figure 1F). The HT eventually closes dorsally (Figure 1G, red arrows in Figure 1G”) and a prominent arterial pole (prospective RV) (Zaffran et al., 2004) becomes visible, completing linear HT formation by ~E8.25 (yellow arrows in Figure 1G”, Figure 1H, see also Video 2).

To assess the overall growth of the forming HT, we measured cTnnT+ tissue volume in segmented z-stacks, at the stages described above (Figure 1I and Supplementary file 1). During the first phase of cardiomyocyte differentiation, the cTnnT signal expands resulting in a cardiac crescent rapidly doubling in volume (Figure 1C’, D’, I). During the subsequent phase of morphogenesis, from cc to open HT stage, growth is less pronounced despite extensive morphological changes (Figure 1E’, F’, I). The Volume of the HT appears to increase again upon addition of the RV primordium to the arterial pole and dorsal HT closure (Figure 1G’, I). HT growth likely reflects an increase in cell number occurring during the formation of the heart tube. Cardiomyocytes are proliferative at this stage (de Boer et al., 2012), and we can indeed observe mitotic figures in the HT (Figure 1J). From this analysis, however, it is unclear how much of the growth observed is due to cardiomyocyte proliferation versus addition of new cardiomyocytes from cardiac progenitor cells located in the splanchnic mesoderm.

**Figure 1.**
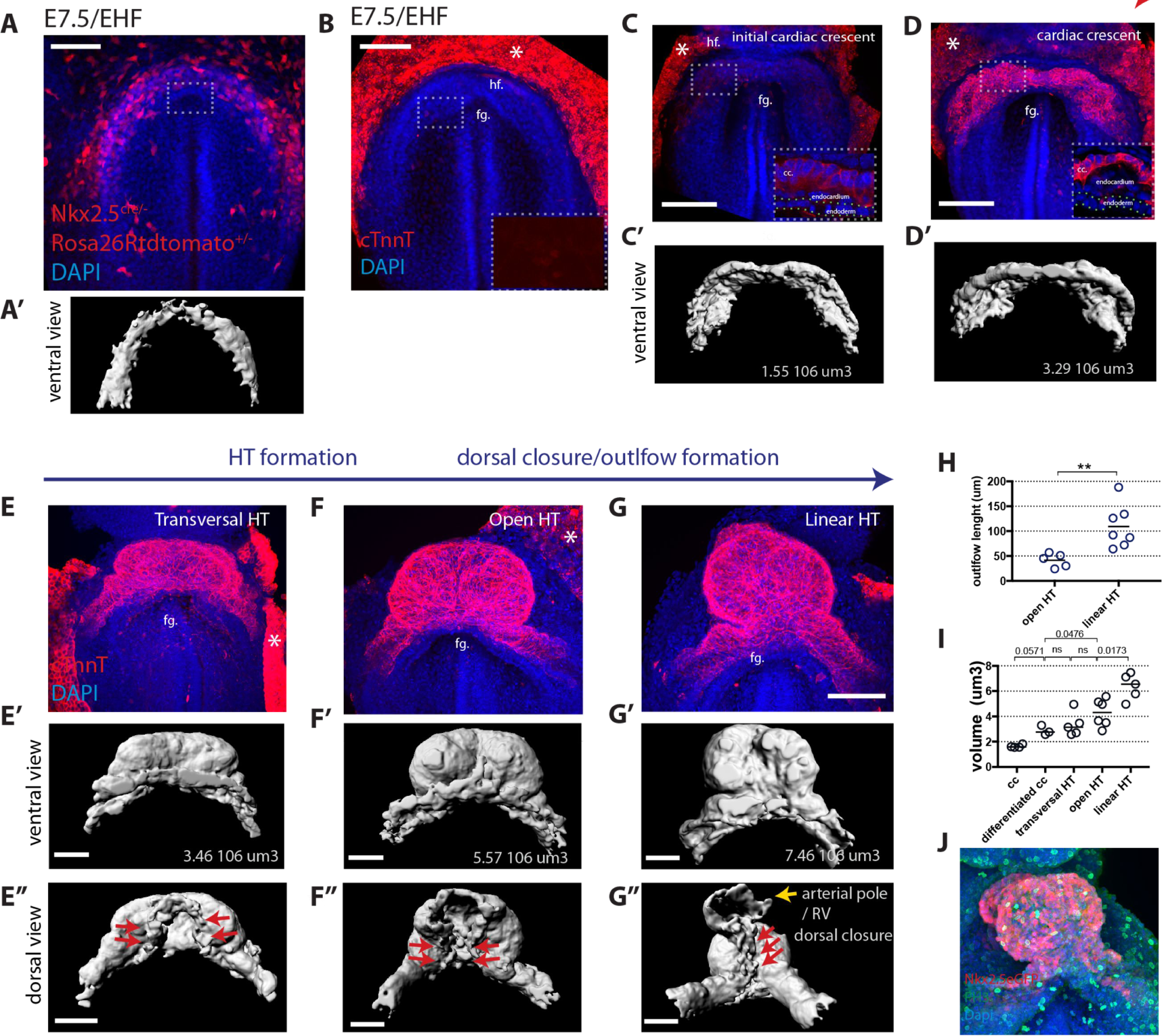
Overview of HT morphogenesis and growth. (A) Frontal view of an *Nkx2.5^cre/+^*; *Rosa26Rtdtomato* embryo at EHF stage. (A’) 3D reconstruction of the tdtomato signal in the cardiogenic area. Signal from tdtomato+ endothelial cells identified by shape was manually masked. See also Video 1. (B-G) Immunostaining for cTnnT (red) and Dapi (blue) showing six consecutive stages during cardiac differentiation (B-D) and HT morphogenesis (E-G). (B) At EHF cTnnt is initially not detectable. (C-D) During early somitogenesis, cTnnT signal becomes detectable in the cc. Insets in (B-D): magnification of single optical sections showing cTnnT localization and cell shape. (C’-G’ and E”-G”) Corresponding 3D renderings from cTnnT signal reconstruction. Red arrows in (E’-G’) highlight the dorsal closure of the HT. Yellow arrow in G” highlights the arterial pole (prospective RV). See also Video 2. (H) Quantification of the arterial pole/RV length in the open HT (41.4 ±14.0μm, n=5) and after dorsal closure (109 ±43.44μm, n=7), mean ± SD, p=0.0025. (I) Quantification of the cardiac volume at the different stages of HT development. (Initial cc: 1.63.106μm3 ± 0.13, n=4, cc: 2.89.106 ± 0.37μm3, n=3, transversal HT: 3.367. 106μm3 ± 0.95, n=5, open HT: 4.29.106μm3 ± 1.08, n=6, linear HT: 6.37. 106μm3 ± 1.01, n=5, mean ± SD). P-values are indicated on the graph. (J) Immunostaining of an *Nkx2.5eGFP* embryo for PH3 (red) and Dapi (blue) at HT stage, showing proliferative cells in the ventricle. Scale bars: 100μm

### 3D analysis of cardiac progenitor differentiation

To visualize the boundary where cardiac progenitors abut differentiating cardiomyocytes during HT morphogenesis, we immunostained *Nkx2.5*^cre/+^ *Rosa26Rtdtomato*^+/−^embryos at the transversal-HT stage with the differentiation marker cTnnT and acquired whole-mount images (Figure 2A). Cre activation of tdtomato was detected in both the FHF and SHF, as well as in the endoderm and endocardium (see optical sections in Figure 2A’, A”, F, F’ and Figure 2-figure supplement 1A) (Stanley et al. 2002). Cardiomyocytes are separated from the endoderm by the endocardium. In contrast, undifferentiated cardiac precursors lie medio-dorsally in direct contact with the endoderm in areas where endocardial cells are not detected (Figure 2A’, A”). This is confirmed by the absence of the endothelial marker CD31 (Figure 2-figure supplement 1A, B, B’). We then rendered in 3D the *Nkx2.5^cre^*-labeled lineages, including both FHF and SHF, and the cTnnT+ tissues, including only the FHF/cc, which allowed visualizing the boundary between SHF and cc at the tissue level (Figure 2A, B, B’, Figure 2-figure supplement 2A’, Video 3 and see material and methods ‐non-cardiac tdtomato signal was removed manually–).

To identify the changes associated to cardiomyocyte differentiation at the cellular level, we labeled single cells with membrane-GFP (see Material and Methods). cTnnT-progenitors have an epithelial-like columnar cell shape, while the differentiated cTnnT+ cardiomyocytes are rounder and have lost the columnar epithelial organization (Figure 2C-E and Figure 2-source data 1). This is reminiscent of the cell shape transition observed in the distal outflow tract (OFT) at later stages of heart development, when SHF progenitor-to-cardiomyocyte differentiation takes place (Francou et al., 2014; Ramsbottom et al., 2014; Sinha et al., 2012). Interestingly, some cells at the boundary zone exhibit weak cTnnT localization and yet show columnar shapes typical of mesodermal cardiac precursors (Figure 2F-F”” and red arrows in Figure 2F”’, F””). Unlike differentiated cardiomyocytes, these cells do not show rounded shapes, and therefore they may represent a transient state between progenitors and differentiated cardiomyocytes; however, the nature of such state cannot be addressed by static analysis. Differentiation of cardiac progenitors is thus accompanied by changes in cell shape and detachment from the endoderm.

We next assessed the expression pattern of the *Nkx2.5eGFP* enhancer reporter line, in which GFP expression is restricted to cardiomyocytes (Lien et al., 1999; Prall et al., 2007; Wu et al., 2006). To characterize this reporter line in detail, we immunostained *Nkx2.5eGFP* embryos against cTnnT, (Figure 3A, B and Figure 3-figure supplement 1A, B) and compared the relative intensities of cTnnT and GFP in manually-segmented single cells (Figure 3C, D Figure 3-figure supplement 2A and Figure 3-source data 1). We found that the GFP level varied linearly with cTnnT level (Figure 3D), although considerable variability of GPF levels was observed within each cTnnT+ and cTnnT-cell population. Scoring of a large number of cells allowed to reproducibly identify the top 50% GFP-expressing cells as positive for cTnnT+ (Figure 3-source data 1). Genetic tracing experiments using the *Nkx2.5^cre/+^ Rosa26Rtdtomato^+/-^* line instead show strong tdtomato level in both the FHF and SHF (Figure 3-figure supplement 3A).

We next characterized the boundary between cardiomyocytes and cardiac precursors in transversal HT stage embryos of the *Nkx2.5eGFP* reporter line. We measured mean fluorescent intensity in manually segmented cells at the boundary zone and found that GFP level and cTnnT signals correlate at the individual-cell level (Figure 3B’, E, Figure 3-figure supplement 2B, C and Figure 3-source data 1). Altogether, these results indicate that the *Nkx2.5-eGFP* reporter is suitable for tracking cardiomyocytes in live-imaging and reliably identifies the top 50% GFP-expressing cells as cTnnT-positive.

**Figure 2.**
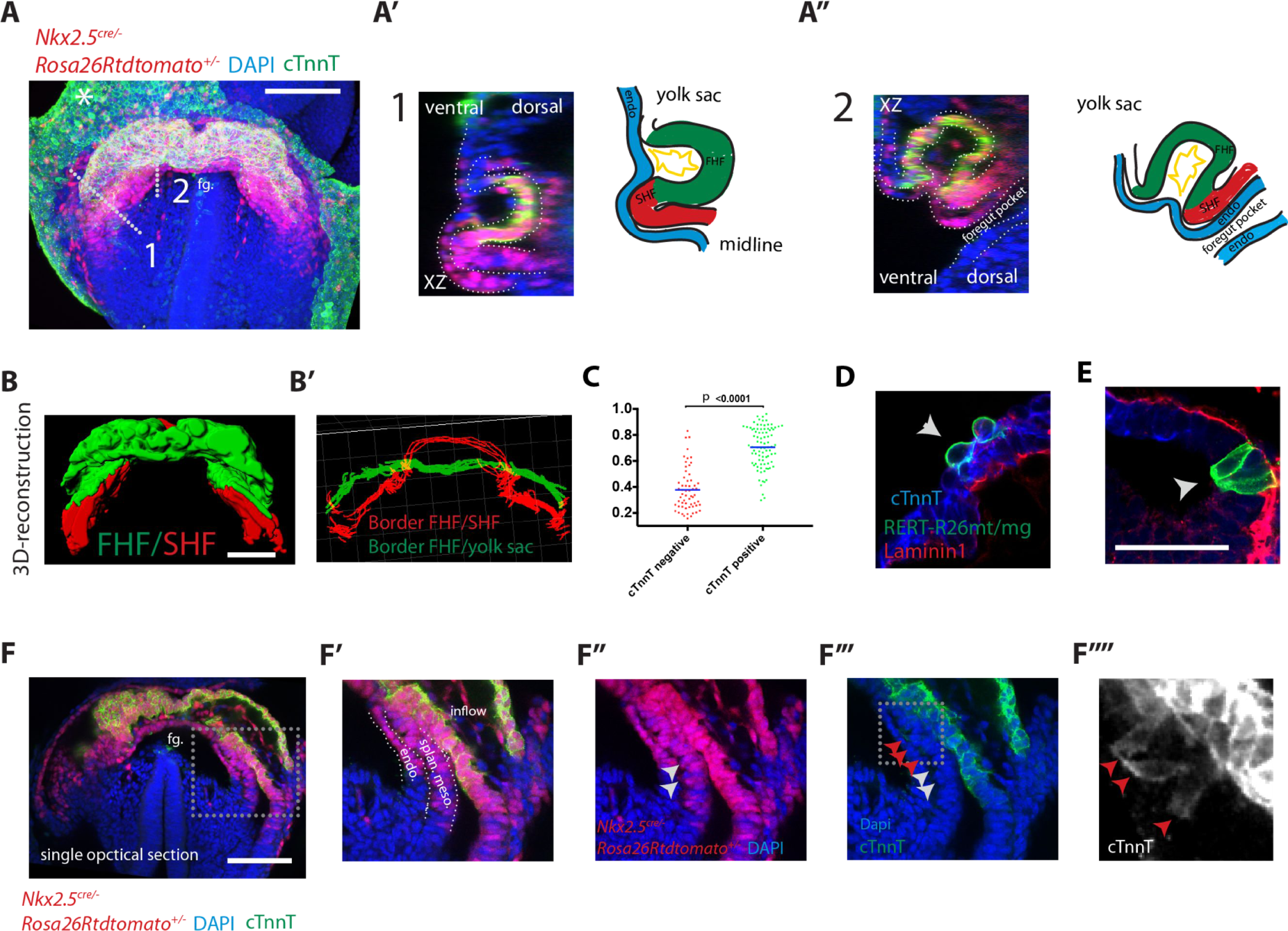
Visualization of the boundary between FHF and SHF. (A-A”) An *Nkx2.5^cre/+^; Rosa26Rtdtomato^+/−^* embryo immunostained for cTnnT (green) and Dapi (blue) showing cells of the Nkx2.5+ lineage populating both the FHF and SHF. (A’ and A”) Cross-sections in xz along the dotted lines 1 and 2 shown in (A), and corresponding schematics highlighting the endoderm (blue), FHF (green), SHF (red) and endocardium (yellow). Note that tdtomato signal is also detected in the endoderm and endocardium. (B) 3D reconstruction of the FHF (green) and SHF (red); (B’) 3D drawing of the border between the FHF and SHF (in red) and the FHF and yolk sack (in green) (based on the embryo shown in (A). For SHF rendering, the tdtomato+ splanchnic mesoderm was depicted. The FHF was rendered using the cTnnT signal. See also Video 3. (C) Quantification of the cell roundness (rnd) index of cardiomyocytes at HT stage on single optical sections. Black bars indicate mean. Rnd index for cTnnT+ (green) cells is 0.71 ± 0.16 and for cTnnT-(red) cells is 0.38 ± 0.16, mean ± SD, exact p value < 0.0001. (D-E) Membrane-GFP labeling of typical cTnnT+ FHF (D) and cTnnT-SHF (E) cells at longitudinal HT stage. The specimen is immunostained for cTnnT (blue), the basement membrane marker Laminin1 (red) and GFP (green). (F) Single optical section of the same embryo shown in (A, B). (F’-F”’) Inset: red arrows point to cells localizing weak cTnnT signal and have columnar cell shape in the SHF. White arrows point to cTnnT-negative SHF cells. splan. meso: splanchnic mesoderm, endo: endoderm, fg: foregut. In all the embryos immunostained for cTnnT, the yolk sack signal is non-specific background (indicated by an asterisks). Scale bars: 100μm except in (D-E): 50μm

**Figure 3.**
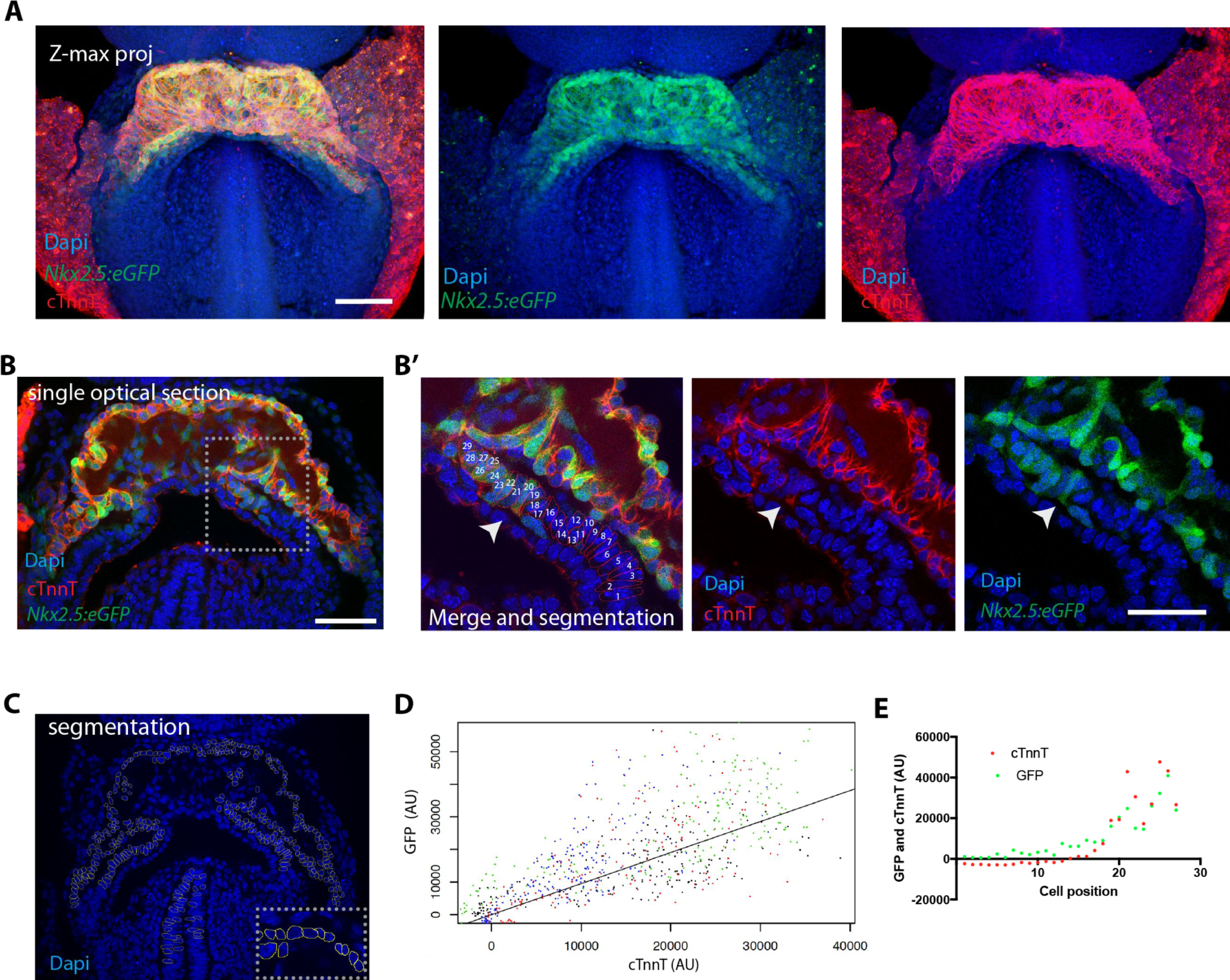
The *Nkx2.5eGFP* transgene is robustly expressed in cTnnT-positive cardiomyocytes. (A) z-maximum projection of an *Nkx2.5eGFP* embryo at transversal HT stage immunostained for cTnnT (red) and Dapi (blue) showing high GFP level in differentiated cardiomyocytes. (B) Single optical section of an *Nkx2.5eGFP* embryo at transversal HT stage immunostained for cTnnT (red) and Dapi (blue). (B’) Inset in (B): arrow points the transition between cTnnT+ and cTnnT-domains, corresponding to FHF and SHF, respectively. Dotted line represents the linescan measured in (D) (C) Example of manual segmentation based on Dapi nuclei in cells located in the splanchnic mesoderm and in the neural tube. (D) Linear mixed-effects model to find the relationship between the background-substracted GFP and cTnnT levels adjusted by embryo (GFP=0.95*cTnnT). (E) GFP and cTnnT mean intensities measured within manually segmented cells along the boundary from SHF to cTnnT-positive FHF (B’) (n=130 cells analysed in 4 independent embryos). Scale bar: 100μm

### 2-photon live-imaging of early cardiac development in the mouse embryo reveals morphogenetic events leading to HT formation

We next established a live-imaging method to dynamically characterize the formation of the HT in the mouse embryo (Figure 4A) (Chen et al., 2014). We adapted a previously reported culture system (Nonaka, 2009; Nonaka et al., 2002) in which the whole mouse embryo is immobilized by inserting the extraembryonic region in a holder (Figure 4C). After culture, embryos showed normal morphology, their hearts were beating and circulation was initiated (Figure 4D, E and Video 4). This culture system in combination with two-photon microscopy enabled the generation of high-resolution 3D+t videos (Figure 3B and see Methods and Materials). Maximum culture time and imaging achieved was 24 hours (Figure 4-figure supplement 1A, n=3). However, typical acquisition times varied in most cases between 3 and 13 hours. The rate of acquisition varied in most cases between 4 and 9 minutes, with some exceptions depending on the specific aim of the recording (see video legends).

**Figure 4.**
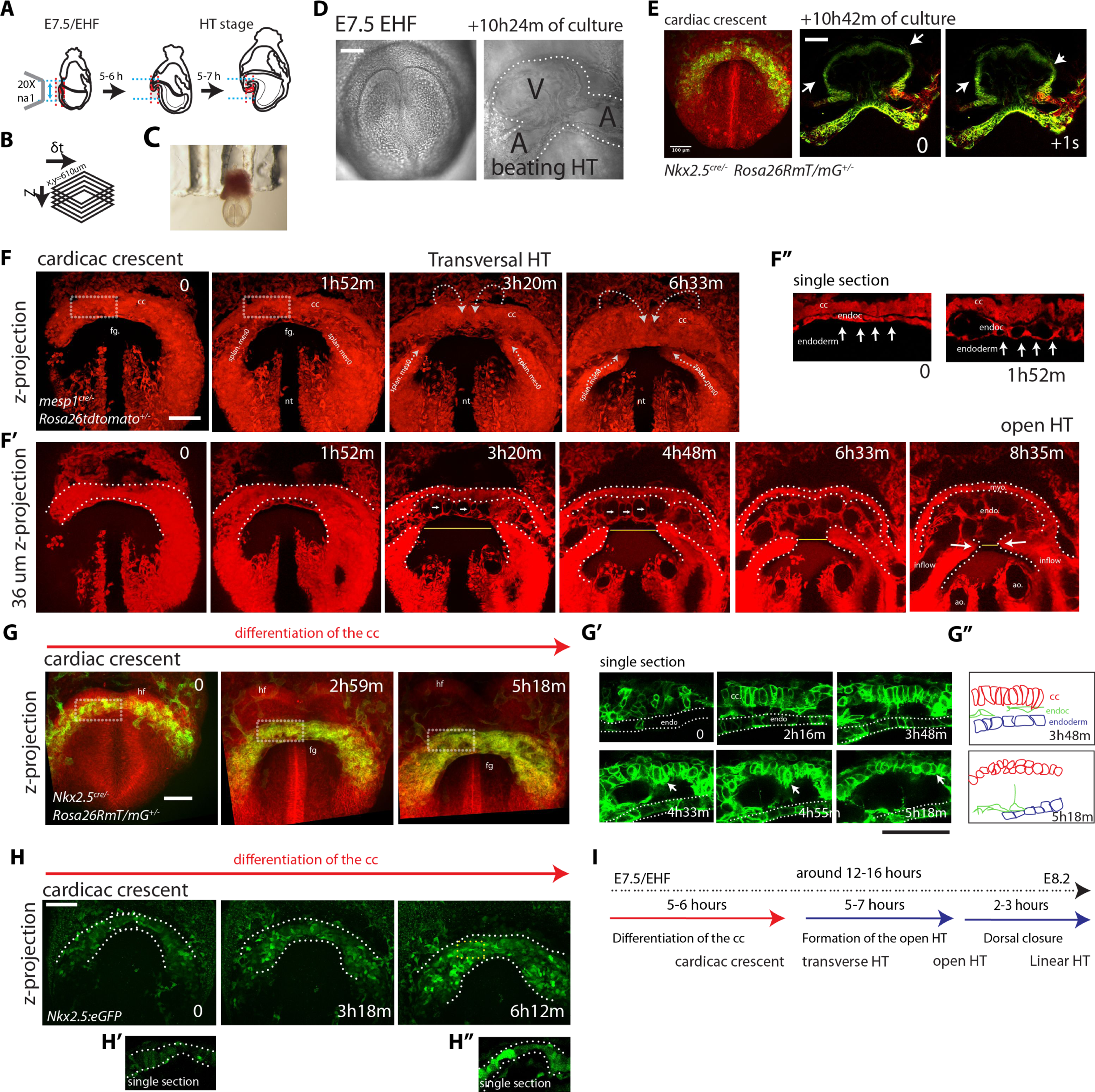
Live-imaging of cardiac differentiation and morphogenesis. (A) Schematic of the set up for imaging live mouse embryos from EHF up to completion of HT formation. (B) Parameters xyzt during live-imaging. (C) An embryonic holder maintains the embryo still during live-imaging. (D-E) After over 10 hours of ex-vivo culture inside the two-photon chamber, the embryo has grown and the cardiac crescent has transformed into a beating heart tube. Arrows in (E) point to the deformation of the heart ventricle due to heart beats. See also Video 4. (F-F”) Time-lapse video sequence of a *Mesp1^cre/+^; Rosa26Rtdtomato^+/−^* embryo ‐reporting anterior mesoderm–, showing the transition from cardiac crescent stage to heart tube stage. Note that at the initial time point, the foregut pocket is already visible. Arrows indicate the major tissue movements visible (folding of the cardiac crescent, medial movement of the splanchnic mesoderm). The differentiation of cardiomyocytes detaching from the endoderm is visible in (F’). White arrows point to the endocardium. The formation of the endocardial lumen in the transversal HT is visible in F’. By 8h35m, the open HT is fully formed and beating regularly. Yellow lines represent distances between the left and right splanchnic mesoderm. Images are z-max projection of 84 sections covering 180μm (F) and 9 sections covering 36μm (F’) acquired every 4μm. See also Videos 5-6. (G-G”) Time-lapse video of an *Nkx2.5^cre/+^; Rosa26RmT/mG^+/−^* embryo during the early stages of cardiac differentiation. Images are z-max projection of 87 sections acquired every 5μm covering 437μm. (G’) shows in a single optical section how progenitors change in cell shape and move away from the endoderm during differentiation towards cardiomyocytes (from inset in (G)). (G”) Cartoon depicting the change in cell shape taking place during cardiac differentiation. See also Video 7. (H) Time-lapse video sequences of *Nkx2.5eGFP* embryos from transversal HT showing low GFP level prior to cardiomyocyte differentiation and increase in GFP intensity level during the stages cardiomyocytes undergo differentiation. (H’ and H”), cardiomyocytes in single magnified optical sections. Images are z-maximum projections of 70 sections acquired every 6μm covering 420μm. See also Videos 10. (I) Estimate of the timing between different heart tube development stages. fg: foregut cc: cardiac crescent, endoc: endocardium, ao: aorta, endo: endoderm. Scale bars: 100μm.

Imaging *Mesp1^cre/+^; Rosa26Rtdtomato^+/-^* embryos allowed tracking the anterior mesoderm (Saga et al., 1996; Saga et al., 1999), including cardiac lineages, from cc stage up to HT stage (Figure 4F, F’ and Videos 5-6). The time-lapse analysis provided insight on the formation of the endocardial lumen (Figure 4F’, F”). The endocardium is initially observed as a bilayer of cells and eventually splits into dorsal and ventral layers, which move apart from each other allowing the lumen to become visible between the layers (see Videos 6 and 7 for confocal views and video 8 for a bright field view). Thin cytoplasmic bridges between the endocardial layers persist and extend as the endocardial layers separate from each other (see white arrows in Figure 4F’ and Video 6). Heartbeat becomes detectable around this stage and circulation in the embryo is then initiated.

Imaging *Nkx2.5*^cre/+^*; Rosa26RmT/mG^+/-^* embryos, in which Cre-recombined cells activate membrane-bound GFP, we could track cells during differentiation, as they transit from a columnar to a round shape and start contracting (Figure 4G-G”, Video 7 and bright-field Video 8). We found that the endocardial lumen started to appear while cc cells still remained columnar (compare time points 2h 13m and 3h 48m in Figure 4G’). Cell rounding is therefore unlikely to initiate the formation of the cardiovascular lumen.

Interestingly, this analysis also showed that the transformation of the cc into the HT involves the antero-medial displacement of the splanchnic mesoderm (white arrows in Figure 4F, F’). This displacement promotes the folding of the cardiac crescent into a hemi-tube by bringing closer the future borders between the HT and the mesocardium (see yellow lines in Figure 4F’ indicating the distance between the borders at different times). These borders coincide with the frontiers between cardiomyocytes and undifferentiated splanchnic mesoderm (see details in Videos 7, 17 and Figure 7A’, A”). Interestingly, the movement of the splanchnic mesoderm is not coherent with the endoderm but a relative displacement is detected between the two layers (Video 9). Estimation of splanchnic mesoderm displacement speed towards the midline from time-lapse analyses (Figure 4F’, and Video 6) indicates a range of average speeds from 12 to 20 µm/hour (15.8 +/-2.4 µm/hour, mean +/− SD, n=3, Figure 4-figure supplement 2 and Figure 4-source data 1). These measurements estimate that midline convergence of the splanchnic mesoderm takes approximately 5-7 hours from the late cc stage until the open HT stage (Figure 4-figure supplement 2).

Our results show the feasibility of live time-lapse analysis of mouse HT formation and reveal that splanchnic mesoderm displacement, at least in part by sliding over the endoderm, is an essential aspect of HT morphogenesis.

### Live tracking of Cardiomyocyte differentiation in individual cells reveals cardiac crescent differentiation dynamics

Next, to specifically track cardiac differentiation, we used the *Nkx2.5eGFP* live reporter. We first studied the general activation pattern of this reporter in live analysis. At E7/bud stage, a faint and scattered GFP signal is detected in proximity to the yolk sack, at the anterior border of the embryo (Figure 5-figure supplement 1A). At neural plate stage, just prior to the ventral folding of the embryo, the GFP signal remains weak but spreads to delineate a crescent in the anterior region of the embryo (Figure 5-figure supplement 1A and Video 10). During about 5-6 hours starting at the EHF stage, the GFP signal increases in intensity, which correlates with the previous observations on cTnnT activation (Figure 4H, Figure 1C and Video 11). From transversal to open HT (5-7 hours) and from the open HT to HT (2-3 hours) the GFP signal remains stable (Figure 5-figure supplement 1B, B’, C, C’ and Video 12). We conclude that an increase in GFP level in *Nkx2.5eGFP* embryos reports cardiomyocyte differentiation. In addition, these results reveal the timing of the main phases of linear HT development; cc differentiation, formation of the open HT and dorsal closure (Figure 4I).

We next sought to track the trajectories and differentiation of individual cardiac precursors within the entire cardiogenic region by 3D+t live imaging. To this end, we used the *Polr2a^CreERT2^* (*RERT*) allele (Guerra et al., 2003), which provides ubiquitous tamoxifen-inducible Cre activity in combination with a *Rosa26Rtdtomato* reporter. We then titrated the tamoxifen dose for a labeling density that would allow single cell tracking during prolonged time-lapse analysis and combined this with the *Nkx2.5eGFP* reporter (see materials and methods). Typically, for each video, we acquired z-slices every 3-5μm achieving a total z-depth of 200μm – with some variations depending on the stage considered- and manually tracked in 3D+t for several hours an initial population of ~50 to ~100 cells per video, which represents around 5-10% of the total number of cells present in the cc (de Boer et al., 2012).

We first tracked cells of the cardiac forming region starting at EHF stage ‐when cardiac precursors are undifferentiated– up to stages in which cardiomyocytes have differentiated in the cc but the transversal HT stage has not been reached yet (Figure 5A and Figure 5-source data 1). At the onset of cardiac differentiation, the cardiac crescent swings ventrally concomitant with foregut pocket formation. During this movement we found that the relative positions of the cardiac progenitors are maintained from the initial stage through the differentiated cc (Video 13). Relative cell positions therefore remain mostly coherent as the embryonic tissues undergo this initial global movement. Differentiation events are detected in some of the tracked cells by cell shape change from columnar to rounded and by the increase in GFP signal (see example in Figure 5B, D and Figure 5-source data 1). In contrast, other tracked cells located in the splanchnic mesoderm remain in contact with the endoderm, retain a columnar shape and show low GFP level throughout the videos (Figure 5C). These cells are likely to be SHF cardiac progenitors and not endocardial cells, which are not present in this area (Figure 2-figure supplement 2A, B, B’). Endocardial cells are instead present in the cardiac crescent and have typical elongated spindle-like shapes (Figure 4F and Figure 5-figure supplement 3A).

**Figure 5.**
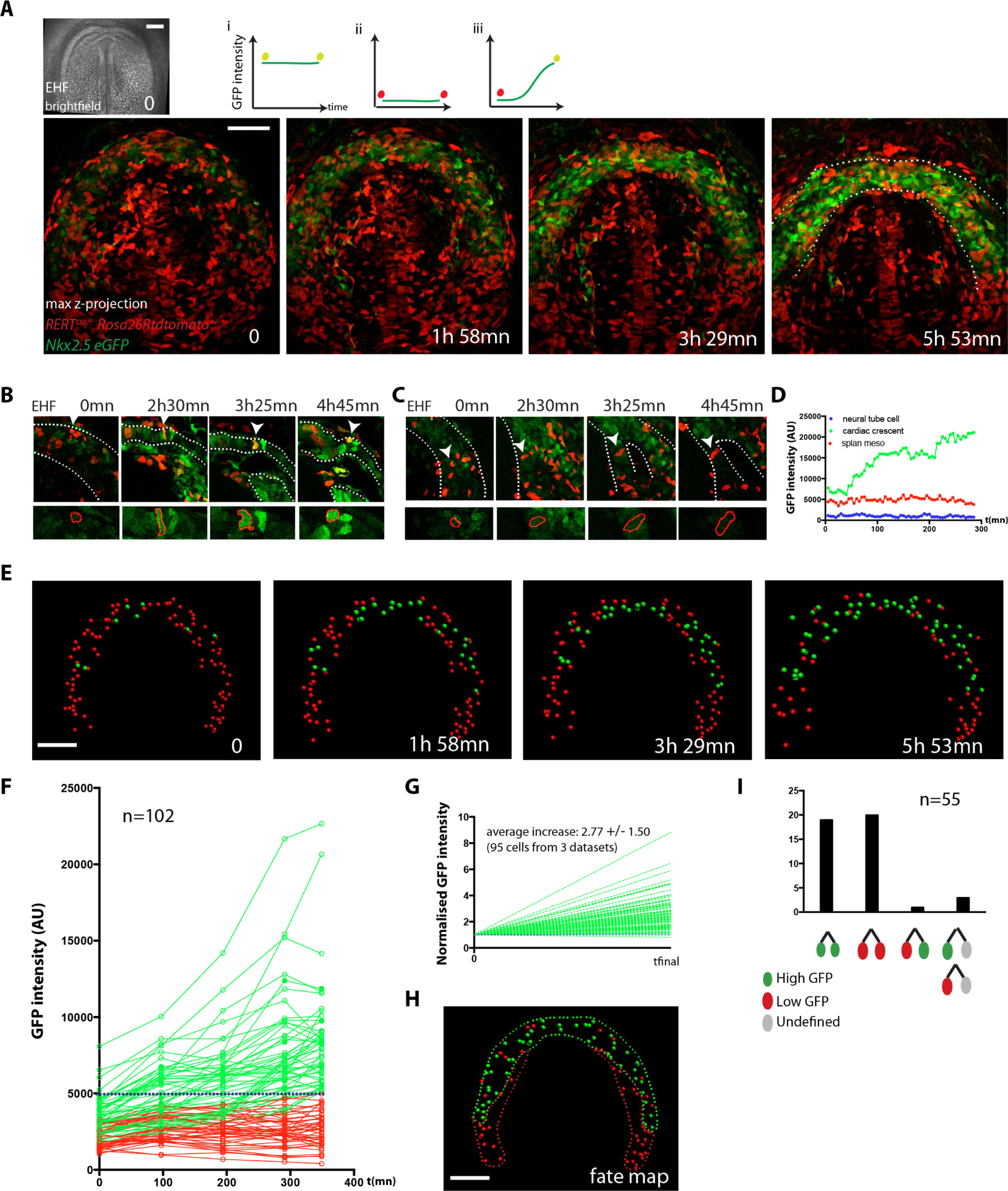
Live-imaging of cardiac differentiation at cellular resolution. (A) Time course of an *Nkx2.5eGFP*; *RERT^cre/+^*; *Rosa26Rtdtomato^+/−^* embryo during stages when cc differentiation takes place-from EHF onwards-. Images are z-maximum projection of 76 sections acquired every 3μm covering 228μm. A bright-field image of the EHF embryo at the initial time point is also shown. (i, ii and iii) Rationale of the expected evolution of GFP expression: since tracks carry information on reporter expression, if a cell acquires a high level of GFP, we predict that it is committed to differentiate. See also Video 12. (B-D) Examples of time-lapse videos (B and C) and quantification of the GFP level (D) increasing in a single cell during differentiation (B) or remaining low in a cell located in the splanchnic mesoderm (C). A neural tube cell was additionally quantified (blue). Images are single optical sections. (E) Time course of individual cells tracked in the video shown in A. Cells with GFP levels above the median intensity value of all cells at the end of the recording are represented as green spheres and cells with lower GFP levels are shown as red spheres. (F) GFP level through time. Blue dotted line: median value (5249 a.u.). GFP level for each tracked cell was measured at five successive time points. Cell divisions are not represented for simplicity. (G) Normalized GFP level showing the relative increase in GFP levels for the GFP-positive cells at the end of the recording. Average increase: 2,77 fold ± 1.50, mean ± SD, n=95 cells from 3 independent videos. (H) The position of cells that differentiate (green) and cells that do not is shown in the cc at the initial time point (EHF stage). (I) The lineage from cardiac precursors share differentiation fate. Lineages of dividing progenitors (n=55 from 3 independent videos) were identified during early stages of cardiac differentiation. Two daughter cells are defined as sharing the same fate if their GFP intensity levels do not differ by more than 1.5 fold and/or both show levels above or below the threshold value defined. Scale bars: 100μm.

Next, in order to establish a fate map of the cardiac forming region at the EHF stage, we tracked back in time the population of cells that showed high GFP intensity (top 50%) at the end of the video (Figure 5F, H and Figure 5-source data 1). According to our previous analysis (Figure 3D), these cells can reliably be assigned to cardiomyocytes. The initial location of this cell population fated to become cc cardiomyocytes delineates a crescent-shaped domain at EHF stage (Figure 5H). Cells retaining lower GFP intensity level throughout the video initially localize posteriorly and medially to this crescent. Most of the cells that have high GFP levels at the final time point show low GFP levels at the initial time point and increase their GFP level over time (Figure 5E-G and Figure 5-source data 1). These results suggest that cardiomyocytes of the cc differentiate during 5-6 hours starting at the EHF stage, which is consistent with the onset of detectable cTnnT at that stage (Figure 1B and Figure 1-figure supplement 2A, A’).

Cells in the cardiac mesoderm do divide during the observation time, so we identified cell division events and tracked the descendant cells. 43% of the tracked cells underwent one division during the 4-5 hour videos. To determine whether cell fate (differentiation versus progenitor) is allocated in the cardiogenic mesoderm at the EHF stage, we tracked GFP levels in dividing cells and their descendants. We found that most sister cell pairs show matched high or low GFP intensity levels at the end of the observation period (38 out of 39, Figure 5I, Figure 5-figure supplement 2A-D and Figure 5-source data 1). This observation suggests that commitment of cardiac precursors to differentiation is already established by the EHF stage and largely transmitted by lineage.

### Cardiomyocyte differentiation is not detected during heart tube morphogenesis

We next studied cardiac differentiation dynamics during subsequent stages when the cc transforms into the HT by extensive morphogenesis. To do so, we tracked cells located in the splanchnic mesoderm and cc in *Nkx2.5-eGFP* embryos at successive periods of around 3 hours covering the 5-7 hours during which the transversal HT transforms into the open HT (Figure 6A, B, E, Video 14 and Figure 4-source data 1). This analysis revealed that the trajectories of cc cells move apart from each other over time as the tissue expands during the transition from the transverse HT to the more spherical open HT. The HT starts to beat during the observation period, especially at the later stages, and therefore, in some cases, cardiomyocyte cell shape appears distorted in single optical sections (Figure 6F), however the GFP level could be determined. As mentioned above, antero-medial movement of the splanchnic mesoderm can be observed concomitant with the transformation of the transversal HT into the open HT (visible also in Video 5 and 12). We found that cells with high GFP level at the initial time points –differentiated cardiomyocytes– retain rather stable GFP levels (green tracks in Figure 6C, D, G and Figure 6-source data 1). In addition, all cells that initially showed low GFP levels did not increase GFP intensity during time-lapse; thus new events of cardiac differentiation were not detectable by this approach (red tracks in Figure 6C, D). Tracking cells for longer periods of time, covering from cc differentiation continuously up to the open HT stage, confirmed that cc differentiation is followed by a period of time in which no differentiation events occur (Figure 6-figure supplement 1, n=19 cells tracked in one embryo).

**Figure 6.**
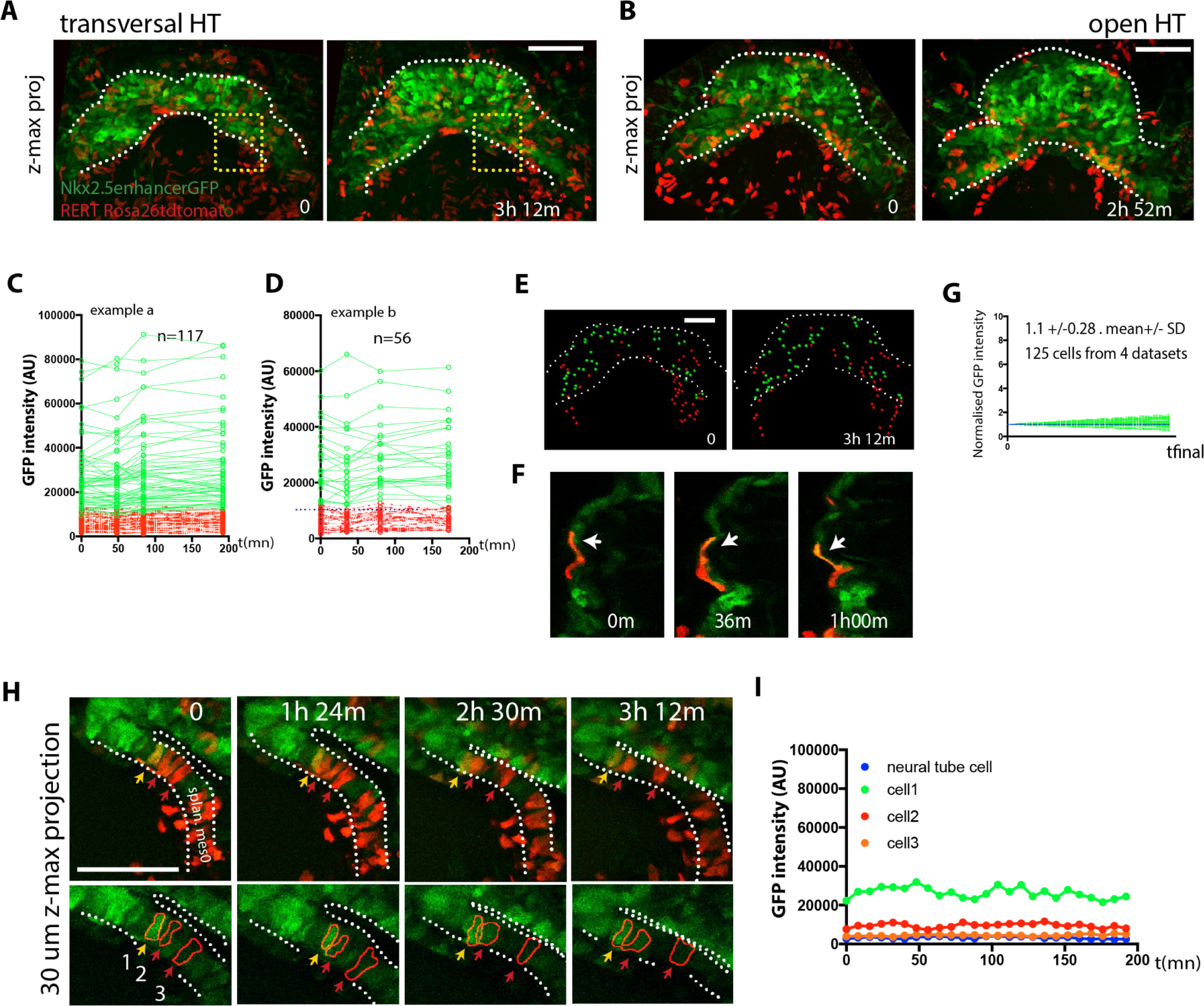
No cardiac differentiation is detected during early HT morphogenesis. (A-B) Initial and final time points of two time-lapse videos of tamoxifen-induced *Nkx2.5eGFP*; *RERT^+/−^*; *Rosa26Rtdtomato^+/−^* embryos covering the transformation of the transversal HT into the open HT. Images are z-maximum projections of 44 sections acquired every 5μm covering 220μm in (A) and of 46 sections acquired every 6μm covering 276μm in (B). See also Video 13. (C, D) GFP levels over time of the cells tracked in (A, B). GFP level for each tracked cell was measured at four successive time points. Green tracks represent cardiomyocytes with GFP intensity above the median intensity value of all the tracked cells at the last time point, while red tracks represent cells with lower GFP level. Blue dotted line represents the median intensity value (10748 a.u. in (C) and 12248a.u. in (D)). Note that when cells divide, only one of the two daughter cells was represented for simplicity. (E) Distribution of the tracked cells from the time-lapse video shown in (A) at the initial and final times of the tracking period. tdtomato+, GFP-cells are represented by red spheres and tdtomato+, GFP+ cells are represented by green spheres. (F) Example of tdtomato+ cardiomyocytes tracked in the beating ventricle. (G) Normalized progression of GFP level in cells classified as GFP+ showing stable GFP levels. Data collected from 125 tracked cells from four independent videos of 2h52m to 3h12m duration. Average increase: 1,1 fold ± 0.28, mean ± SD. (H) Magnification extracted from the video in A (yellow inset) of tdtomato+ cells in the boundary zone between the undifferentiated and differentiated cells. (I) Quantification of GFP level through time of the three segmented cells shown in (H). Cells show stable GFP levels and do not differentiate. Images are z-maximum projections of 6 sections acquired every 5μm covering 30μm. Scale bars: 100μm.

To confirm the absence of detectable cardiac differentiation events during this period, we next focused on the live analysis of cells located at the boundary between cardiomyocytes and undifferentiated splanchnic mesoderm. As expected, we observed GFP-low cells located adjacent to GFP-high cells in the boundary zone (Figure 6H and Video 14). Those cells retain stable GFP levels throughout the tracking time and did not increase their GFP level (Figure 6I, boundary imaged 20 times in different locations and in 6 independent embryos). Importantly, they retain a columnar shape typical of weak cTnnT+ and cTnnT-cells located at the boundary zone (see Figure 2F” and Figure 3B’). Although these cells migrate antero-medially relative to the underlying more static endodermal cells (see endodermal cell highlighted by the red arrow from t=69m in video 9), they do not contribute to the HT during the observation period. We confirmed this observation in longer time-lapse videos spanning 7 hours that covered the whole transition from transversal to open HT stage. Again, progenitors strictly respected the boundary with the HT throughout the entire time-lapse video (Figure 7A-A”’ and Video 15, boundary imaged 5 times in distinct locations and in 2 independent embryos). All together, these data suggest that during the transformation of the cc into the dorsally open HT no cardiomyocytes are added to the HT from the SHF. These observations suggest two distinct phases of early HT formation: a first phase of differentiation of the FHF into the cc, lasting around 5 hours, and a second phase of HT morphogenesis in which the SHF progenitors remain undifferentiated, lasting around 5-7 hours. During this second phase, extensive remodeling of the cardiac crescent is concomitant with the antero-medial splanchnic mesoderm displacement.

**Figure 7.**
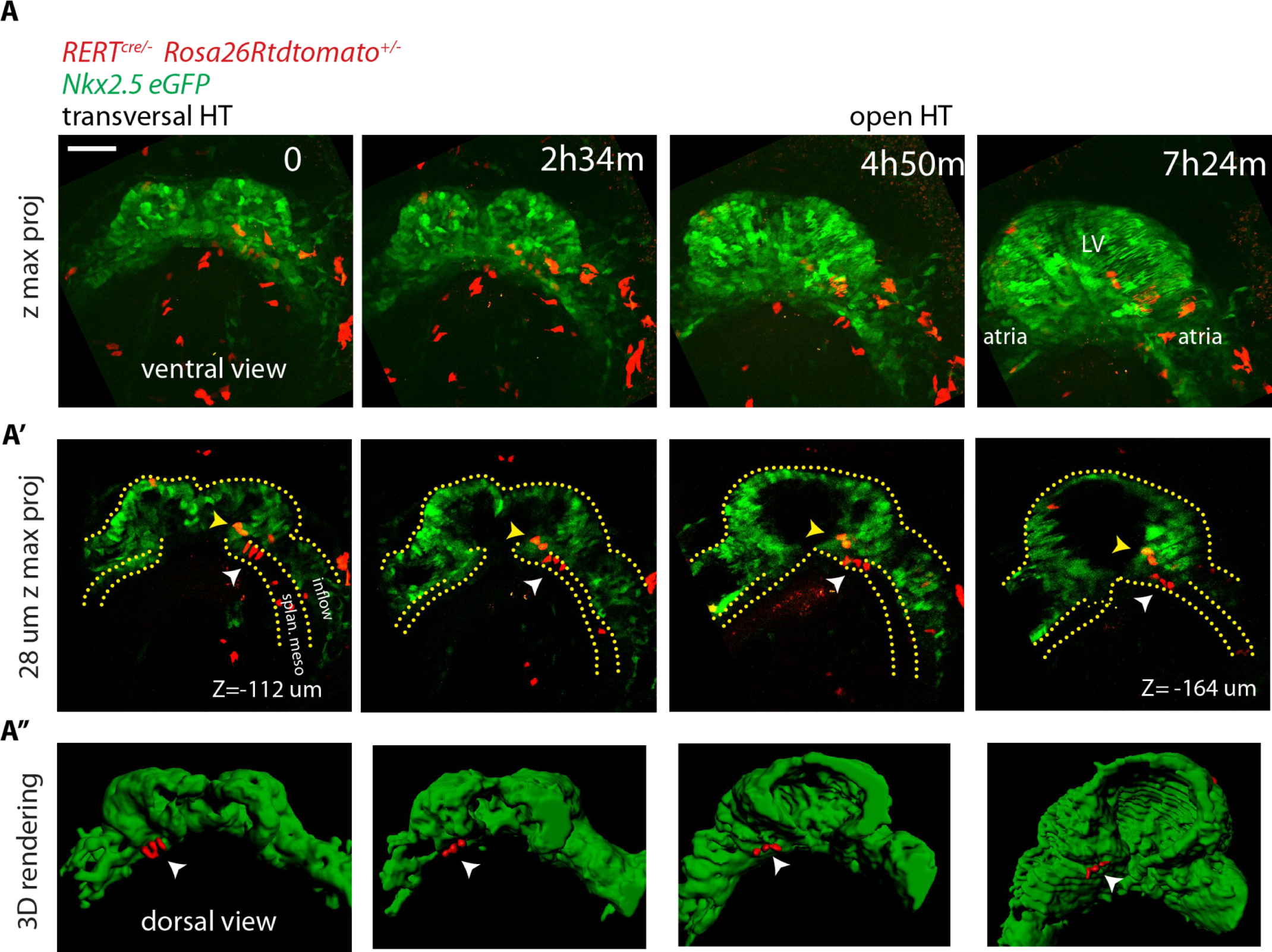
No cardiac differentiation is detected during early HT morphogenesis. (A-A”) Time-lapse video of a tamoxifen-induced *RERT^+/-^; Rosa26Rtdtomato^+/−^; Nkx2.5eGFP* embryo during the stages at which the transversal HT transforms into an open HT, showing cells located in the splanchnic mesoderm (white arrows in A’ and A”), respecting the boundary with the HT (yellow arrows in J’ show two cells located in the HT). Lower doses of tamoxifen were injected in order to label only a very small proportion of cells in red. Images are z-maximum projection of 74 sections (A) acquired every 4μm covering 296μm and (A’) 7 sections covering 28μm. (A”) 3D reconstruction based of the *Nkx2.5eGFP* signal (green) and tdtomato+ cells located in the splanchnic mesoderm. See also Video 14. Scale bars: 100μm. Splan. meso: splanchnic mesoderm.

### Imaging the Isl1-expressing cell lineage confirms absence of cardiac progenitor differentiation during early heart tube morphogenesis

The LIM domain transcription factor Islet1 (Isl1) is a cardiac progenitor marker. Its expression is transient in the precursors of the cc, while it remains expressed in SHF progenitors for an extended period (Brade et al., 2007; Prall et al., 2007, Yuan and Schoenwolf, 2000). Cells of the *Isl1*-expressing lineage detected with Cre reporters therefore contribute only scarcely to the cc, while extensively to the SHF and its derivatives (Cai et al., 2003; Ma et al., 2008). To test these observations in live imaging, we combined *Nkx2.5eGFP* with tracing of the *Isl1* cell lineage using the *Isl1^cre^* driver and the *Rosa26Rtdtomato* reporter. We found that tdtomato labeling is first detectable in scarce isolated cells of the GFP+ cc during the period when the cc swings ventrally and differentiates (from t=2h 36m to t=4h in Figure 8A, B). In contrast, a dense tdtomato labeling appears in the GFP-low cells of the splanchnic mesoderm as the cardiac crescent fully differentiates (from t=2h24m to t=4h in Figure 8A and C and Video 16). tdtomato is as well detected in the endoderm and endocardium (not shown). Consistently with previous reports (Cai et al., 2003), *Isl1^cre^*-induced recombination detected in live analysis is thus low in the cc and complete in the SHF.

**Figure 8.**
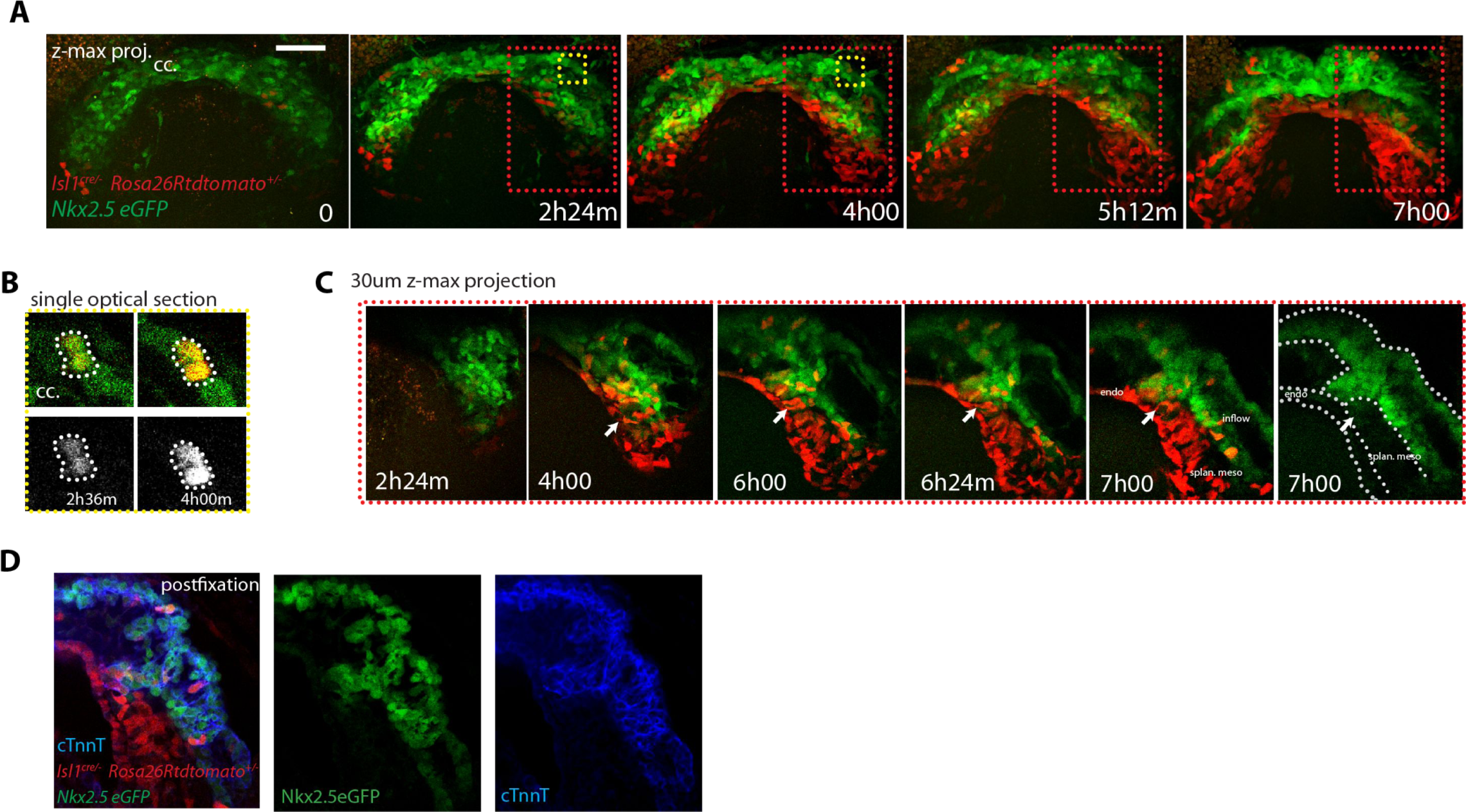
SHF Isl1-expressing cells do not contribute to the early open-heart tube. (A) Time-lapse video sequences of an *Isl1^cre/+^; Rosa26Rtdtomato^+/−^; Nkx2.5eGFP* embryo showing that recombination driven by *Islet1^cre^* of the *Rosa26* locus is complete in the splanchnic mesoderm and scarce in the FHF (n=3 videos from cc up to open HT stage). (B) Inset from A (yellow frame). Two cells of the Isl1+ lineage located in the cc increase their tdtomato level over time. (C) Inset from A (Red frame); increase of the tdtomato intensity in the splanchnic mesoderm over time. Arrow shows the boundary between SHF and FHF and points to a single cell in the SHF that retains a low level of GFP and a columnar shape. Note that tdtomato signal is also detectable in the endoderm. Images are z-maximum projection of 45 sections acquired every 5μm and covering 225μm in (A) and of 6 sections covering 30μm in (C). See also Videos 15-16. (D) Same embryo as in (A) post-fixed and immunostained against cTnnT after live-imaging, showing that the red cells located in the splanchnic mesoderm are undifferentiated. Scale bars: 100μm.

Once the cc is formed, if cells of the SHF would continuously differentiate, then regions of the forming heart tube contributed by the SHF precursors should appear densely co-labeled with both GFP and tdtomato. Live imaging shows instead that tdtomato+ cells establish a boundary with GFP+ cells, confirming no signs of differentiation of SHF precursors during the observation period (Figure 8A, C and Videos 17). The live imaging, however, did not allow to unambiguously identify all cells located deep inside the live tissue at the final stages recorded. The arterial pole in particular is located deep in the embryo. It is therefore challenging to accurately track cardiac differentiation by the increase of GFP levels there (Figure 8-figure supplement 1A-A”, from around 200μm depth, see next section). To overcome these limitations, we fixed and immunostained embryos against cTnnT after completion of the live-imaging experiments, and imaged them by 3D confocal microscopy. No solid domains containing double-labeled cells were detected, indicating that progenitors located in the SHF did not undergo differentiation in the boundary zone from cc to open HT stage (Figure 8D). These results are consistent with the single-cell tracking analysis and confirm that the SHF does not differentiate during linear HT morphogenesis.

### Cardiac differentiation is detected during the late stages of HT development

We next wanted to determine when cardiac progenitors located in the SHF start to differentiate. In order to address the timing of SHF contribution to the arterial pole, we next fixed and optically cleared *Nkx2.5eGFP*; *Ils1*^cre/+^; *Rosa26Rtdtomato*^+/−^ embryos at different stages from cc up to heart looping (n=10) and assessed the appearance of GFP and tdtomato double-positive domains in the HT. In agreement with our previous observations, we found that SHF cells do not differentiate up to the open HT stage, when the dorsal seam of the heart is still open. In contrast, massive appearance of solid domains of double positive cells is observed subsequently in the fully closed HT, reinforcing our previous interpretation (Figure 9A, B and Videos 19-20). At this stage, the primordium of the RV has been added at the arterial pole (Zaffran et al., 2004, Laugwitz et al., 2005, Moretti et al., 2006) and is fully composed of double-positive cells. The dorsal seam of the HT is also densely populated by double-positive cells, indicating a contribution of precursors from the splanchnic mesoderm to the cardiomyocyte population that finalizes the dorsal closure of the linear HT.

**Figure 9.**
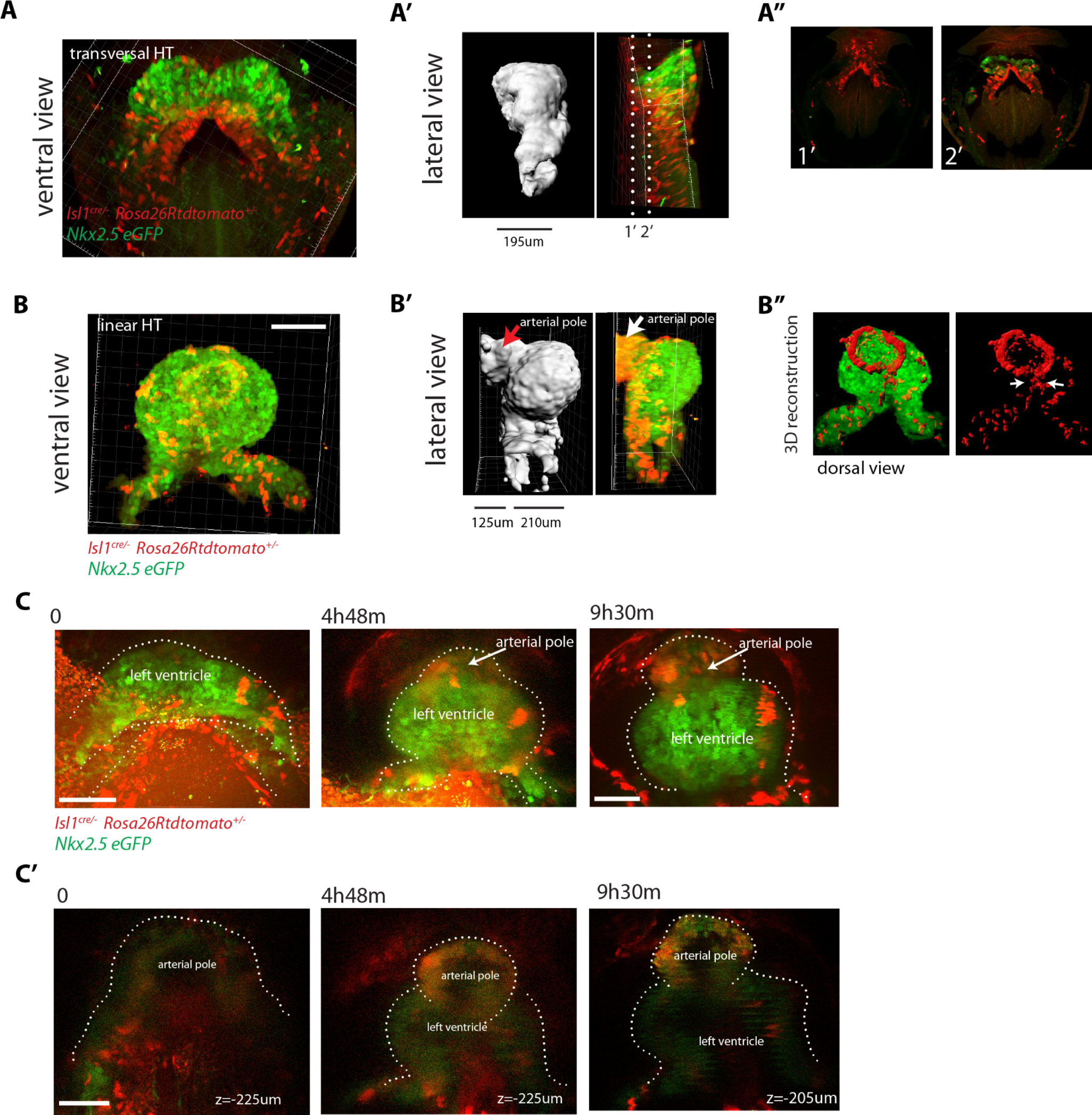
SHF Isl1-expressing cells contribute to the arterial pole during late linear HT morphogenesis. (A, B) *Isl1^cre/+^; Rosa26Rtdtomato^+/−^; Nkx2.5eGFP* embryos showing no contribution of the Isl1 lineage to the transversal HT (A-A”) but contributing robustly to the arterial pole and dorsal aspect of the linear HT (B-B”). The presence of double-positive cells in these areas reveals the differentiation of Isl1 lineage cells into cardiomyocytes (A, B). Images are 74 and 134 optical sections acquired every 2.5μm and covering 195μm and 335μm, respectively. In (B, B’) only the double-positive cells located in the linear HT are shown, while the tdtomato+ progenitors, located outside the linear HT are not shown. Lateral views are shown in (A’, B’) including 3D reconstructions. (A”) Cross-sections in xy along the dotted lines shown in (A’). Dorsal views are shown in (B”) including the 3D reconstruction of the tdtomato+ cells located in the linear HT. White arrows show the dorsal (mesocardial) regions of the linear HT. (C) Snapshots of an *Ils1^cre/+^; Rosa26Rtdtomato^+/−^; Nkx2.5eGFP* embryo at different time points during the third phase showing that the tdtomato+ SHF cells contribute to the arterial pole of the HT. Images are 52 optical sections acquired every 5μm and covering 260μm. The embryo was imaged continuously every 13m for the first 6h of culture and then imaged once again at time point 9h30m. Note that at the last time point, the laser was increased at maximum power in order to reveal more clearly the red labeled cells at the arterial pole. The contrast had to be enhanced as well. (C’) Images at time points 0, 4h48m and 9h30m. Images at time points 0 and 4h48m are z-maximum projection of 5 sections covering 25 μm. Image at 9h30m is a single optical section. at: arterial pole. endo: endoderm, splan meso: splanchnic mesoderm. Scale bars: 100μm.

In order to capture the initiation of SHF contribution to the HT by live imaging, we first focused on the venous pole, as this region is more directly exposed than the rest of the HT and therefore more suitable for live-imaging. In videos that captured the transition from the open HT to the closed linear HT, some cells at the border between the FHF and SHF maintain low GFP levels during the first part of the recording and upregulate GFP to the level of the cardiomyocyte population as the HT forms (Figure 8-figure supplement 2). We next aimed to live-track the activation of SHF differentiation at the arterial pole. Because of the imaging limitations in this area, quantitative analysis of the GFP signal was not possible and we instead used the qualitative detection of SHF cells addition to the HT. To achieve this, we imaged an *Nkx2.5eGFP*; *Ils1*^cre/+^; *Rosa26Rtdtomato*^+/−^ embryo at successive time points throughout the transition from open to linear HT and tracked tdtomato+ cells incorporation to the HT at the arterial pole. This study further confirmed that SHF differentiation at the arterial pole is initiated when the HT is about to close dorsally but not before (Figure 9C). Thus, SHF cells do not differentiate during the 5-7 hour period when morphogenesis of the open HT takes place, but coordinately start differentiation at different regions of the HT during dorsal closure. Interestingly, these regions do not only include arterial and venous poles but also the dorsal seam of the HT.

## DISCUSSION

Here we established a whole-embryo live-imaging method based on two-photon microscopy that allows whole-tissue tracking at cellular resolution. By combining various genetic tracing tools, we labeled progenitor and differentiated cardiomyocytes and performed 3D cell tracking over time combined with 3D reconstruction of the HT at multiple stages. We report three distinct temporal phases of HT formation (Figure 10). During the first phase, the cc differentiates rapidly and morphogenesis, in terms of changes in the relative position of cells, is minimal. During the second phase, differentiation is not detected and morphogenetic remodeling gives rise to a dorsally open HT. During the third phase, cardiac precursor recruitment and differentiation resumes, contributing to the formation of the RV and the dorsal closure of the HT. Our results support the early establishment of distinct FHF and SHF cell populations and show that the morphogenetic changes that transform the cc into a HT largely take place in the absence of cardiac precursor differentiation. These observations indicate tissue-level coordination of differentiation and morphogenesis during early cardiogenesis in the mouse.

**Figure 10.**
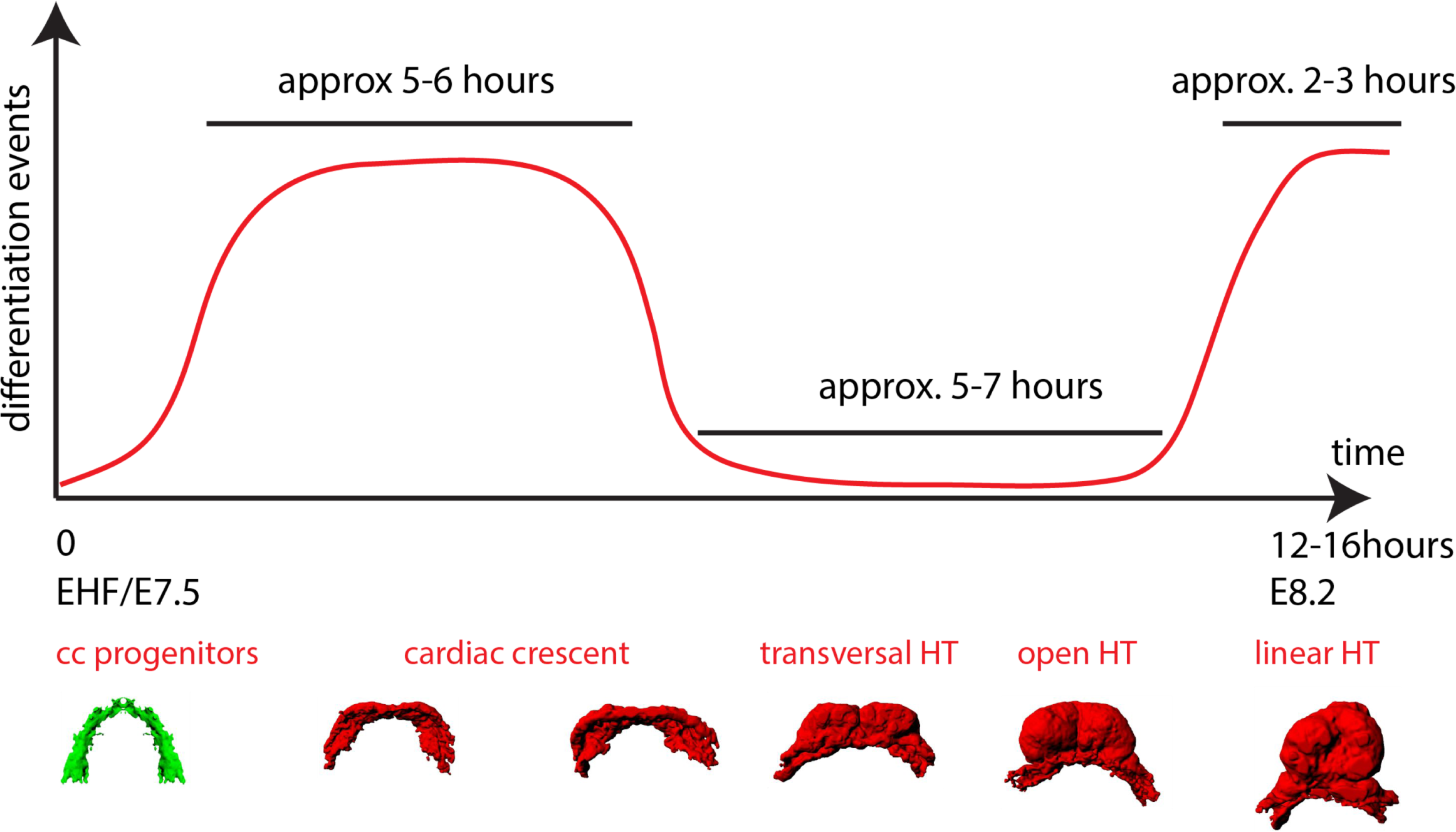
A model of cardiomyocyte differentiation dynamics during early heart development. We propose that two distinct phases of cardiomyocyte differentiation take place during early heart development. At EHF stage the cc differentiates and starts folding. Cardiomyocytes round up and become contractile while the cardiac progenitors located in the splanchnic mesoderm remain undifferentiated. Subsequently, the cc. undergoes further morphogenesis to transform into a HT, initially open dorsally and no cardiomyocyte differentiation is detected during this transformation. Finally, cardiac differentiation resumes contributing new cardiomyocytes from the splanchnic mesoderm to the arterial pole (prospective RV) and the dorsal closure of the HT.

The series of 3D reconstructions from fixed embryos was important to establish a reference staging of HT formation. This allowed us to accurately stage embryos in live experiments based on morphology and it will also be useful in the future for gene expression mapping and accurate phenotypic analysis of mutant embryos. The tissue growth pattern observed in static 3D reconstructions was insightful to suggest variability in growth rates during different phases of HT formation. Growth of the differentiated cardiac tissue is relatively paused when the cc undergoes morphogenesis to form the open HT during the second phase. This is consistent with previous studies in mouse, chick and human models showing that proliferation drops in the differentiated myocardium of the forming HT, while proliferation remains high in the splanchnic mesoderm (Gert van den Berg et al. 2009, de Boer, B.A, 2012, Sizarov et al. 2010). Our live analysis further showed that SHF cells do not contribute to the forming HT during the differentiation pause, which correlates with the growth rate reduction during this phase. This period coincides as well with the onset of cardiac contractility in the embryo (Tyser et al. 2016).

Following the phase of differentiation pause, growth of the HT is reinitiated by incorporation of new cells as the HT closes dorsally and the RV precursors are added at the arterial pole during the third phase. During this third phase, similarities were found in the differentiation dynamics of SHF precursors and splanchnic precursors contributing to the dorsal regions of the linear HT. The dorsal aspect of the linear HT gives rise to the inner curvature of the looped heart, which has an important contribution to non-chamber myocardium, including atrio-ventricular canal and parts of the conduction system (Christoffels et al., 2000). Our results suggest that the late recruitment of progenitors to the dorsal HT could contribute to differences between inner curvature cardiomyocytes and the rest of the heart tube.

While the live-imaging experiments were essential for the identification of the 5-7 hour hiatus between FHF and SHF differentiation, live imaging of the arterial pole during the SHF differentiation phase was challenging and was complemented by 3D reconstructions based on fixed and optically cleared embryos. These experiments confirmed the pause in differentiation during open HT formation and its reactivation during linear HT closure. In the future, it will be interesting to explore whether novel non-toxic index-matching media compatible with embryo viability (Boothe et al. 2017) may alleviate the limitations for deep cardiac imaging during late HT formation.

Regarding the specification of FHF and SHF populations, previous prospective clonal analyses showed that these lineages diverge around gastrulation (Devine et al., 2014; Lescroart et al., 2014). In agreement with this, our tracking of cell lineages in the cardiac forming region shows that sister cells share fates to either the cardiac crescent or the SHF. In addition, the fact that cells contributing to the SHF do not differentiate during the period when the cardiac crescent transforms into the primitive heart tube may contribute to the establishment of the sharp boundary observed between left and right ventricles later in development (Devine et al., 2014). Further studies will be required to assess how this temporal pause of cardiac differentiation is regulated. The molecular analyses of early FHF and SHF precursors suggest that intrinsic molecular differences between the two lineages appear around or shortly after gastrulation (Lescroart et al. 2014). These intrinsic differences may contribute to the regulation of the two distinct differentiation schedules described here. These studies and our observations, however, cannot discriminate whether this lineage allocation results from intrinsic differences between these lineages or it is due to their exposure to position-specific environments, especially as in our studies sister cells remain close neighbors. Environmental cues thus could also control the sequential differentiation of FHF and SHF precursors. The Wnt and BMP pathways are well known regulators of cardiac differentiation (Ai et al., 2007; Jain et al., 2015; Klaus et al., 2007; Kwon et al., 2007; Marvin et al., 2001; Qyang et al., 2007; Tirosh-Finkel et al., 2010; Ueno et al., 2007) and specific mechanisms affecting these pathways could be operating during the formation the HT, whereby the differentiation pathways could be temporally restrained. Finally, the endoderm may also play a key role in mediating FHF differentiation. Indeed cardiac mesoderm differentiation is affected in the absence of Sox17 a transcription factor required for maintenance of the definitive endoderm‐ and this is accompanied by the formation of a morphologically abnormal HT (Pfister et al. 2011). It remains however unclear whether this effect is secondary to an initial defect in foregut development that is observed in this mutant.

A recent study reported spontaneous calcium transients propagating laterally thought the cardiac crescent (Tyser et al. 2016). Left/Right (L/R) asymmetry therefore exists within the cardiac crescent, prior to any detectable cardiac contraction, and BMP/SMAD1 signaling in the lateral plate mesoderm may be involved in this asymmetry (Furtado et al. 2008). In our studies, however, we did not detect any L/R difference in cTnnT expression by whole-mount immunodetection or Nkx2.5GFP activation by live analysis in the cc.

Regarding the possible conservation of the temporal differentiation sequence described here, elegant experiments in zebrafish using a cardiac myosin light chain reporter line and a Kaede photo-conversion assay, addressed the temporal order of cardiac differentiation in live embryos (de Pater et al., 2009; Liu and Stainier, 2012). Two distinct phases of cardiomyocyte differentiation were also observed. During a first phase, cardiomyocytes were recruited first into the ventricle and atria at the venous pole. During a second phase, cardiomyocyte differentiation was observed at the arterial pole of the HT. These pulse-chase experiments however do not address whether cardiac differentiation is continuous or includes a differentiation-paused phase, so further studies will be required to establish the conservation of the observations made here for the mouse embryo.

Finally, an important question to address is the functional relevance of the three distinct phases described here; more specifically, what is the role of the observed differentiation pause. An interesting possibility is that this pause would be functionally related to the extensive morphogenetic events that transform the cardiac crescent into the HT. Our study demonstrates coordinated splanchnic mesoderm movements during this phase. These movements involve a very active antero-medial displacement of the splanchnic mesoderm surrounding the cc -mostly fated to the SHF-. This behavior of the splanchnic mesoderm appears essential for transforming the cc into a dorsally closed HT. Importantly, this displacement involves the sliding of the splanchnic layer over the endoderm, suggesting an active role of SHF precursors in this morphogenetic movement. Such displacement of the splanchnic mesoderm over the endoderm during cardiogenesis had been suggested by classical time-lapse studies in the chick embryo (de Haan, 1963) and it is tempting to speculate that a similar phenomenon contributes to the incorporation of SHF cells to both poles of the HT at later stages of HT development (van den Berg et al. 2009; Kelly RG, et al. 2001; Zaffran et al, 2004). These observations suggest that SHF cells do not represent just a reservoir of cardiac precursors but play a morphogenetic role essential for heart tube formation. The main consequence of the displacement of the splanchnic mesoderm layer over the endoderm is the medial convergence of the left and right frontiers between the cc and SHF to form the dorsal mesocardium and close the HT. An important consequence of the differentiation pause described here is the stability of the cc/SHF frontiers during these morphogenetic movements. This stability prevents further spreading of the differentiation wave from the cc into the SHF/splanchnic mesoderm, which could interfere the ability of the latter to efficiently displace over the endoderm. We therefore hypothesize that the stability of the cc/SHF frontiers –and thus the differentiation pause-would be essential to allow the effective displacement of SHF/splanchnic mesoderm and elicit HT formation. The temporal allocation of the morphogenetic phase after cc differentiation allows the formation of the HT while simultaneously providing an incipient cardiac function essential for the organization of embryonic circulation. This hypothesis poses a functional basis for the alternation of differentiation and morphogenesis phases during HT formation.

Our study applies whole-embryo live analysis of cardiac development at tissue level and with cellular resolution. We expect that extending this experimental approach will allow to further uncover unexpected and novel mechanisms of organogenesis. While limited attention had been paid so far to the temporal dynamics of differentiation during embryonic development, this is an essential aspect of organogenesis (Gogendeau et al., 2015; Parchem et al., 2015; Yang et al., 2015). Here we show the relevance of differentiation timing regulation during heart tube formation and its coordination with morphogenesis at the tissue level. Further understanding of the molecular and cellular mechanisms underlying these phenomena will help us expanding pools of cardiac progenitors in vitro or directing them towards differentiation.

## MATERIALS AND METHODS

### Mouse Strains

Mouse alleles used in the manuscript are listed including bibliographic references and allele identities at the “Mouse Genome Informatics” data base (MGI, http://www.informatics.jax.org/). *Mesp1^cre^* (Saga et al., 1999, MGI:2176467), *Isl1^cre^* (Cai et al., 2003, MGI:3623159), *Nkx2.5^cre^* (Stanley et al., 2002, MGI:2448972), *Rosa26Rtdtomato* (Madisen et al., 2010, MGI:3809524), *Rosa26RmTmG* (Muzumdar et al., 2007, MGI:3716464), *Nkx2.5eGFP* (Wu et al., 2006, MGI:5788423), *Polr2a^CreERT2^* (*RERT*) (Guerra et al., 2003, MGI:3772332) and *C57BL/6* (Charles River). Mice were genotyped as previously described. All animal procedures were approved by the CNIC Animal Experimentation Ethics Committee, by the Community of Madrid (Ref. PROEX 220/15) and conformed to EU Directive 2010/63EU and Recommendation 2007/526/EC regarding the protection of animals used for experimental and other scientific purposes, enforced in Spanish law under Real Decreto 1201/2005.

### Immunostaining and Imaging

Embryos dissected in Dulbecco’s modified Eagle’s medium (DMEM, Invitrogen) were fixed overnight in 2% PFA at 4C, then permeabilized in PBST (PBS containing 0.1% Triton X-100) and blocked (5% goat serum). Embryos were incubated overnight at 4ºC with antibodies diluted in PBST: mouse anti-cTnnT (1:250, MS-295 Thermo Scientific), rabbit anti-PH3 (1:250, 06-570 Millipore), CDC31 (553370 BD Pharmingen™ clone MEC 13.3), SMA (C6198 Sigma) and rabbit anti-Laminin1 (1:500, Sigma, L9393). After washing in freshly prepared PBST at 4ºC, embryos were incubated with secondary antibodies (Molecular Probes, A21121, A21141, A11035) coupled to 488, 549 or 649 fluorophores as required at 1:250 and DAPI at 1:500 (Molecular Probes, D3571) overnight at 4ºC. Before imaging, embryos were washed in PBST at room temperature and cleared with focus clear (Cell Explorer, FC-101) to enhance the transparency of the embryo. Confocal images were obtained on a SP8 Leica confocal microscope with a 20X oil objective (0.7 NA) at a 1024 × 1024 pixels dimension with a z-step of 2-4μm. Embryos were systematically imaged throughout the entire heart tube from top to bottom.

### 3D Reconstruction and Volumetric Measurement

For 3D rendering, fluorescent signal in confocal z-stacks was first segmented by setting intensity thresholds using the trainable Weka segmentation tool plugin available in Fiji (Arganda-Carreras et al., 2017; Schindelin et al., 2012). The resulting z-stacks were then corrected manually on a slide-by-slide basis to eliminate segmentation mistakes. In case of the cTnnT immunofluorescence images (Figure 1), background signal from the yolk sack was manually masked. The volume of the cTnnT positive myocardium was then computed by multiplying the total segmented area by the z-stack interval using a custom Fiji macro. In the *Nkx2.5^cre/+^; Rosa26tdtomato^+/-^* and *Nkx2.5eGFP* embryos, fluorophore signal present in the endothelium, endocardium and endoderm cells was manually masked prior to segmentation (Figure 1A, Figure 1-figure supplement 1, Figure 2B, Figure 2-figure supplement 2A,A’ and, Figure 6A”). For 3D visualization of the 3D segmented image stacks, Imaris software (Bitplane) was used.

### Embryo Culture and Two-photon Live-Imaging

Embryos were dissected at E7.5 in pre-equilibrated DMEM supplemented with 10% foetal bovine serum, 25 mM HEPES-NaOH (pH 7.2), penicillin (50μml21) and streptomycin (50mgml21). Embryos were staged on the basis of morphological criteria (supplementary Figure 1) (Downs and Davies, 1993; Kirstie A. Lawson, 2016), and those between the bud and early somitogenesis stages were used for culture and time-lapse imaging. To track the early phase of cardiac differentiation and subsequent phases of morphogenesis, we used embryos at EHF to transversal HT stage. Embryos were cultured in 50% fresh rat serum, 48% DMEM without phenol red, 1% N-2 neuronal growth supplement (100X, Invitrogen 17502-048) and 1% B-27 supplement (50X Thermo Fisher Scientist 17504044) filter sterilised through a 0.2 mm filter. To hold embryos in position during time-lapse acquisition, we made special plastic holders with holes of different diameters (0.5-3 mm) to ensure a good fit of embryos similarly to the traps developed by Nonaka et al. (Nonaka, 2009; Nonaka et al., 2002). Embryos were mounted with their anterior side facing up. To avoid evaporation, the medium was covered with mineral oil (Sigma-Aldrich; M8410). Before starting the time-lapse acquisition, embryos were first pre-cultured for at least 2 hours in the microscopy culture set up. The morphology of the embryo was then carefully monitored and if the embryos appeared unhealthy or rotate and move, they were discarded, otherwise, time-lapse acquisition was performed. For the acquisition, we used the Zeiss LSM780 equipped with a 5% CO2 incubator and a heating chamber maintaining 37ºC. The objective lens used was a 20X(NA=1) dipping objective, which allowed a long working distance for imaging mouse embryos and tissues. A MaiTai laser line at 1000 nm was used for 2-channel 2-photon imaging. Acquisition was done using Zen software (Zeiss). Typical image settings were: output power: 250mW, pixel dwell time: 7μs, line averaging: 2 and image dimension: 610x610μm (1024x1024 pixels). To maximize the chance of covering the entire heart tube during long-term time lapse videos, we allowed 150-200μm of free space between the objective and the embryo at the beginning of the recording.

### Cell labeling, 3D tracking and GFP Intensity Measurement

For labeling single cells, tamoxifen was administered by oral gavage (2-4 mg/mL) in *RERT;Rosa26R-tdtomato* (cell tracking) or *RERT;Rosa26RmTmG* mice (cell shape study) at E7. Cell shape measurements were done on single cells, imaged in mosaic-labeled, fixed and immunostained embryos and analyzed with Fiji software (Figure 2D, E). Tracking of tdtomato-labeled cells was done on single cells located within the cardiogenic mesoderm -excluding endothelial, pericardial and endodermal cells‐ and their GFP intensity was measured over time. To track cells manually in 4D stacks, the MTrackJ Fiji plugin (Meijering et al., 2012) was used. A local square cursor (25x25 pixels) on the cell of interest snaps according to a bright centroid feature on a slice-by-slice basis. Only tracks lasting for the entire length of the video were kept. When an ambiguity arises in the tracking between consecutive time points, the track was discarded. Tracks split at cell divisions. A cell division event is normally clearly distinguishable over at least 2 time points. In case one of the two daughter cells is not tractable, the other daughter cell is still tracked. Each track is assigned an ID number and excel files with all the tracks coordinates in x, y, z and t was generated. Coordinates of each track were converted into 8-bit 4D images using a custom Fiji macro in which each cell was represented by a sphere of specific pixel intensity, from 1 to 255, while pixels corresponding to background were set to zero. The 4D images were then opened with Imaris to perform visualization of the 3D trajectories of each cell using the “spots” tool, where each object were identified according to pixel intensity. GFP intensity measurement is performed by segmentation of cell shape. A Gaussian filter whose radius is adjusted to the typical size of a cell was first applied, followed by a Laplacian filter. The resulting 32 bits image was next converted to a mask by thresholding. When objects touched each other, a watershed on the binary mask and manual corrections was applied. Each segmented cell was checked and tracked manually for accuracy. In Figure 3B’ nuclei segmentation was performed manually. The mean GFP signal intensity of the segmented objects was then measured using the “analyze particle” tool in Fiji. To quantify the GFP level of tracked cells through time, four to five successive time points were arbitrarily chosen in each video (Figure 5F, Figure 6C, D and Figure 5-figure supplement 1C) except in Figure 5D and Figure 6I, where GFP intensity level was measured in every time point. Background intensities were measured in neural tube cells, which are known to be negative for GFP and cTnnT. Tables containing ID number of tracked cells and GFP intensities were generated and plotted using Prism statistical software.

### Statistical Analysis

For comparisons of two groups, a Man-Whitney U-test was used using Prism statistical software. To find a correlation between GFP and cTnnT levels of 0.8 with an alpha-level f 0.05 and a power of 0.2 at least 10 cells per embryo were required (Figure 3D and Figure 3-figure supplement 2C). Many more cells were computed for each experiment. The linear fit was done using “lm” function from R statistical software (https://www.r-project.org/). To calculate the average speed of splanchnic mesoderm displacement, the shortest distance between the left and right splanchnic mesoderm was calculated in two z-level, using the measure tool in Fiji, and the variation in this distance by time unit was divided by 2, to determine the speed of movement of each sliding side of the splanchnic mesoderm.

## Acknowledgments

We thank the CNIC Microscopy unit for help with the live confocal analysis, Fatima Sanchez Cabo for help with the statistical analyses and Florencia Cavodeassi, Miguel Manzanares, Briane Laruy and members of the Torres lab for helpful comments on the manuscript. This work was supported by grants BFU2015-71519-P, BFU2015-70193-REDT and RD16/0011/0019 (ISCIII) from the Spanish Ministry of Economy, Industry and Competitiveness (MEIC). KI was supported by a Human Frontiers Science Program (LT000609/2015) and EMBO (ATL1275-2014) postdoctoral fellowships. The CNIC is supported by the Spanish MEIC and the Pro CNIC Foundation, and is a Severo Ochoa Center of Excellence (MINECO award SEV-2015-0505). The authors declare no conflicts of interest.

## FIGURE LEGENDS

**Figure 1-figure supplement 1.**
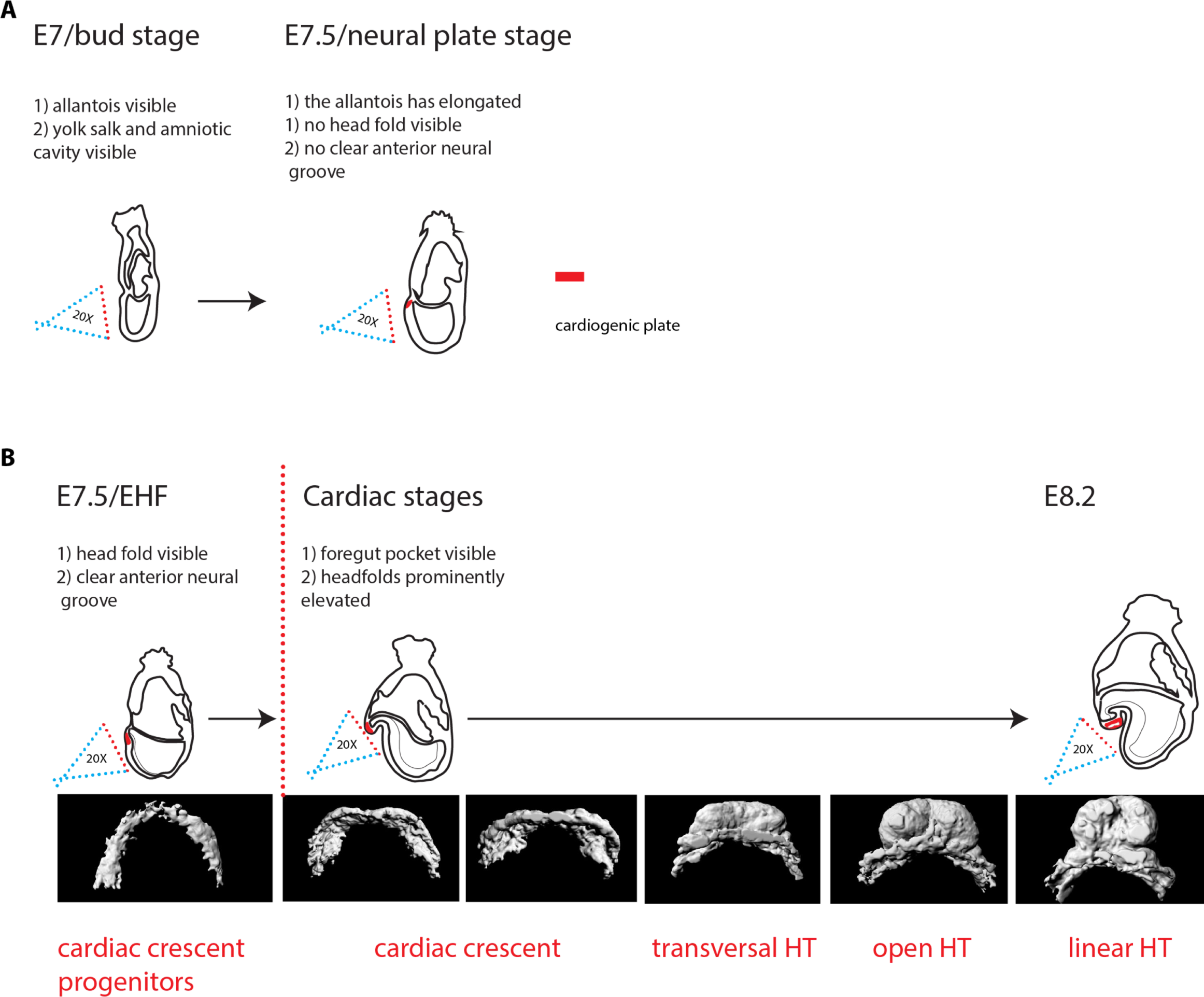
Criteria for embryo staging. We determined the developmental stage of embryos based on morphological landmarks (Downs and Davies, 1993; Kirstie A. Lawson, 2016). (A) At bud stage/E7, a small allantois bud is visible. At neural plate stage/E7.5, the allantois is larger and projects into the yolk sack cavity. The anterior neuroectoderm is enlarged. (B) At EHF/E7.5, the neural plate starts to form the head fold. During somitogenesis, a clearly visible foregut pocket appears and the head folds are located dorsally and anteriorly to the cardiac primordium. The cc has differentiated. It transforms successively into the transversal HT, the open HT and the linear HT closed dorsally and with a prominent arterial pole/RV. Scale bars: 100μm.

**Figure 1-figure supplement 2.**
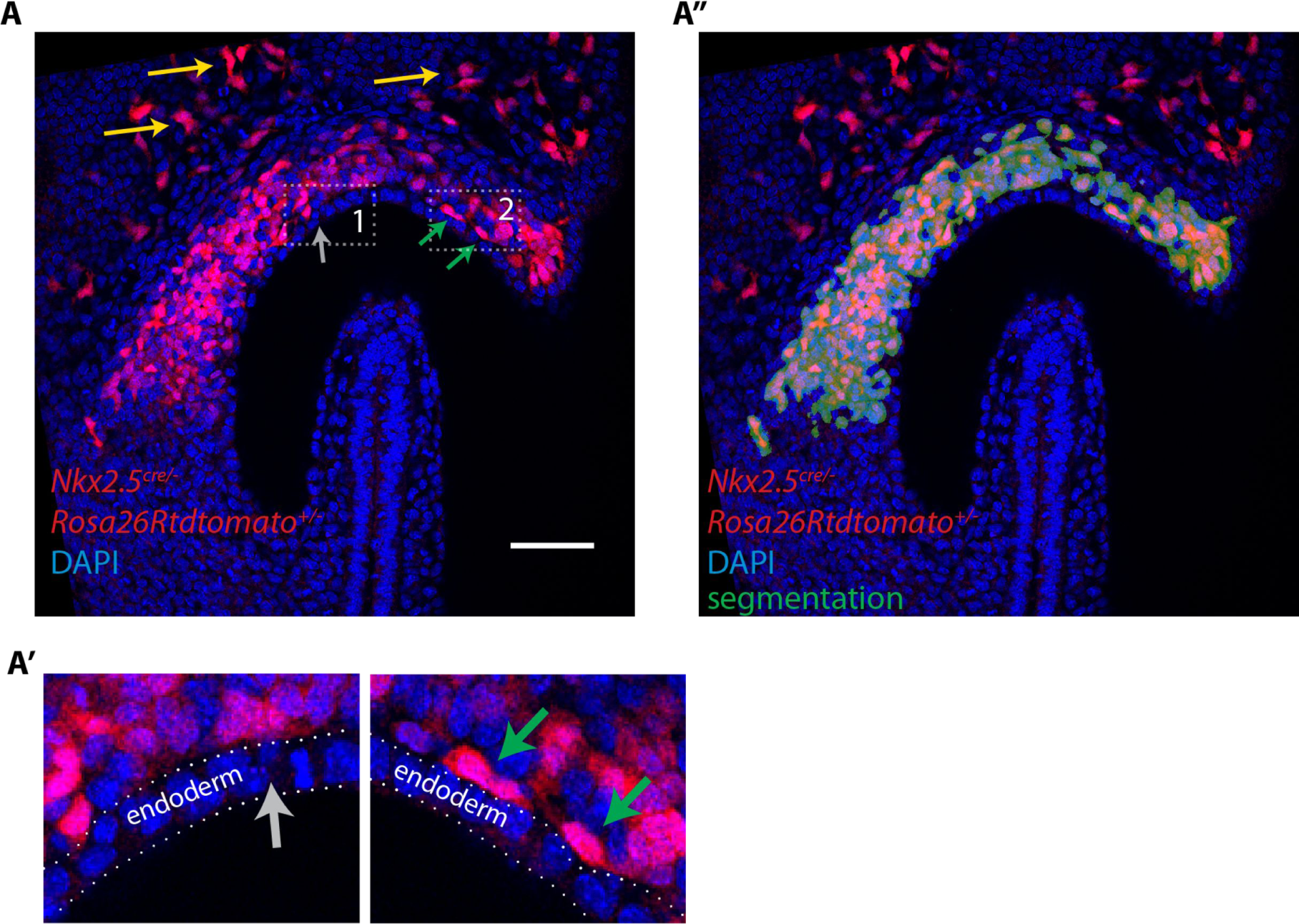
Lineage tracing using the *Nkx2.5^cre^* driver shows a contribution of labeled cells to the cardiogenic regions but not to the endoderm at EHF stage. (A, A’) *Nkx2.5^cre/+^; Rosa26Rtdtomato^+/−^* embryo at EHF stage (same embryo as in Figure 1A. Here a single optical section is shown). Tdtomato labeling is detected in the cardiac progenitors, endothelial (yellow arrow) and endocardial cells (green arrows). No labeling is detected in the endoderm (gray arrow). (A”) Segmentation of cardiac crescent progenitors (shown in dark green) used for the 3D reconstruction shown in Figure 1A’ overlaid with the raw image and. Scale bars: 100μm

**Figure 1-figure supplement 3.**
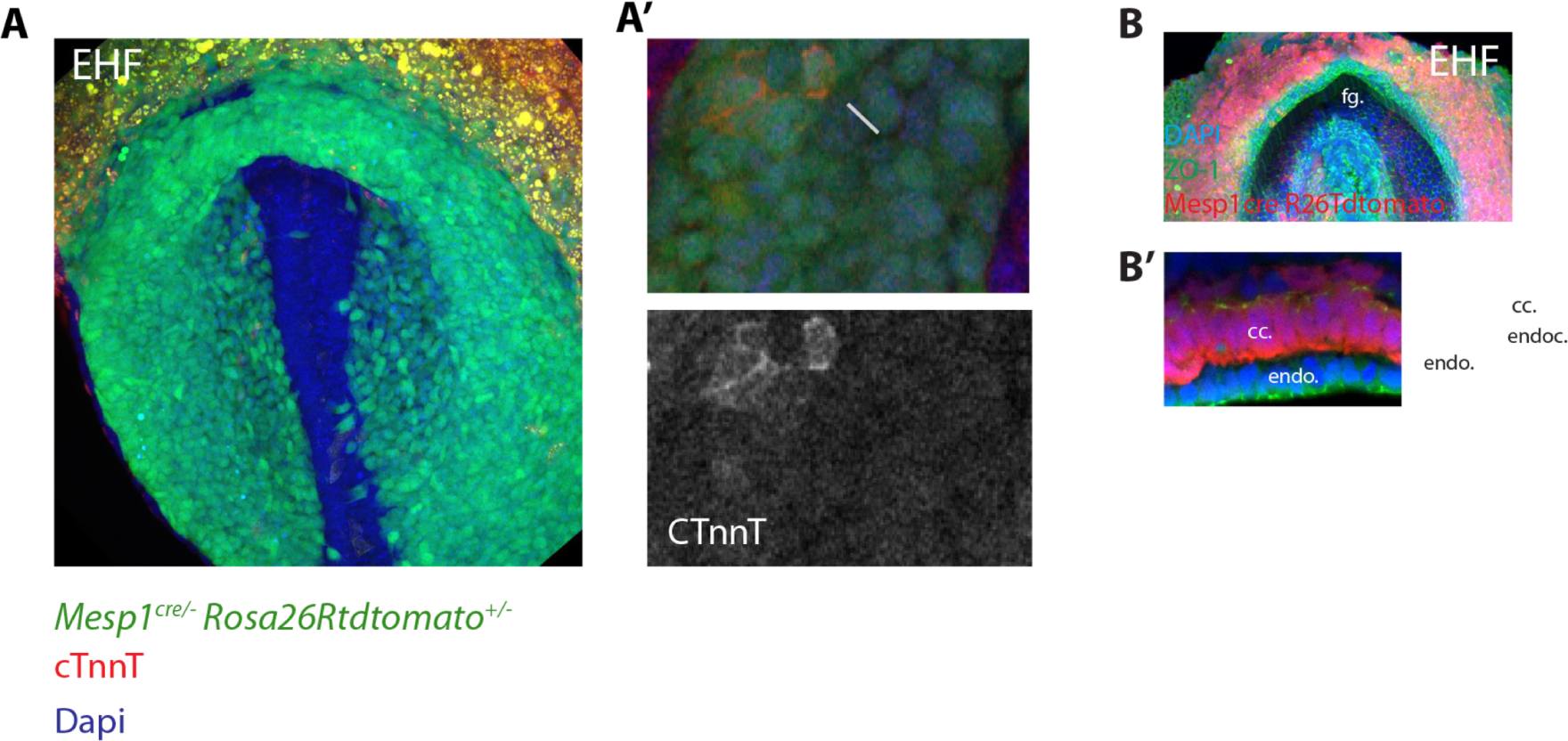
Faint cTnnT signal starts to be detected at EHF stage in apico-basally polarized cc cells. (A) A *Mesp1^cre/+^; Rosa26Rtdtomato^+/−^* embryo at EHF stage showing mesodermal labeling (green)-immunostaining against cTnnT (red) and Dapi (blue). Note that the embryo is at a slightly more advanced developmental stage than those shown in (Figure 1B) because the foregut pocket is more invaginated. (A’) White inset in (A). Faint cTnnT signal (red) in few mesodermal cells (arrows) can be detected. (B-B’) *Mesp1^cre/+^; Rosa26Rtdtomato^+/−^* embryo showing the mesoderm in red and immunostained against the tight junction component zona-occludens-1 (ZO-1) (green) and Dapi (blue). The cc cells (as seen in transversal sections) have an AB polarized epithelial morphology. fg: foregut, endo: endoderm, endoc: endocardium. Scale bars: 150μm.

Figure Source Data 1

Source data for Figure 1H and I

**Figure 2-figure supplement 1.**
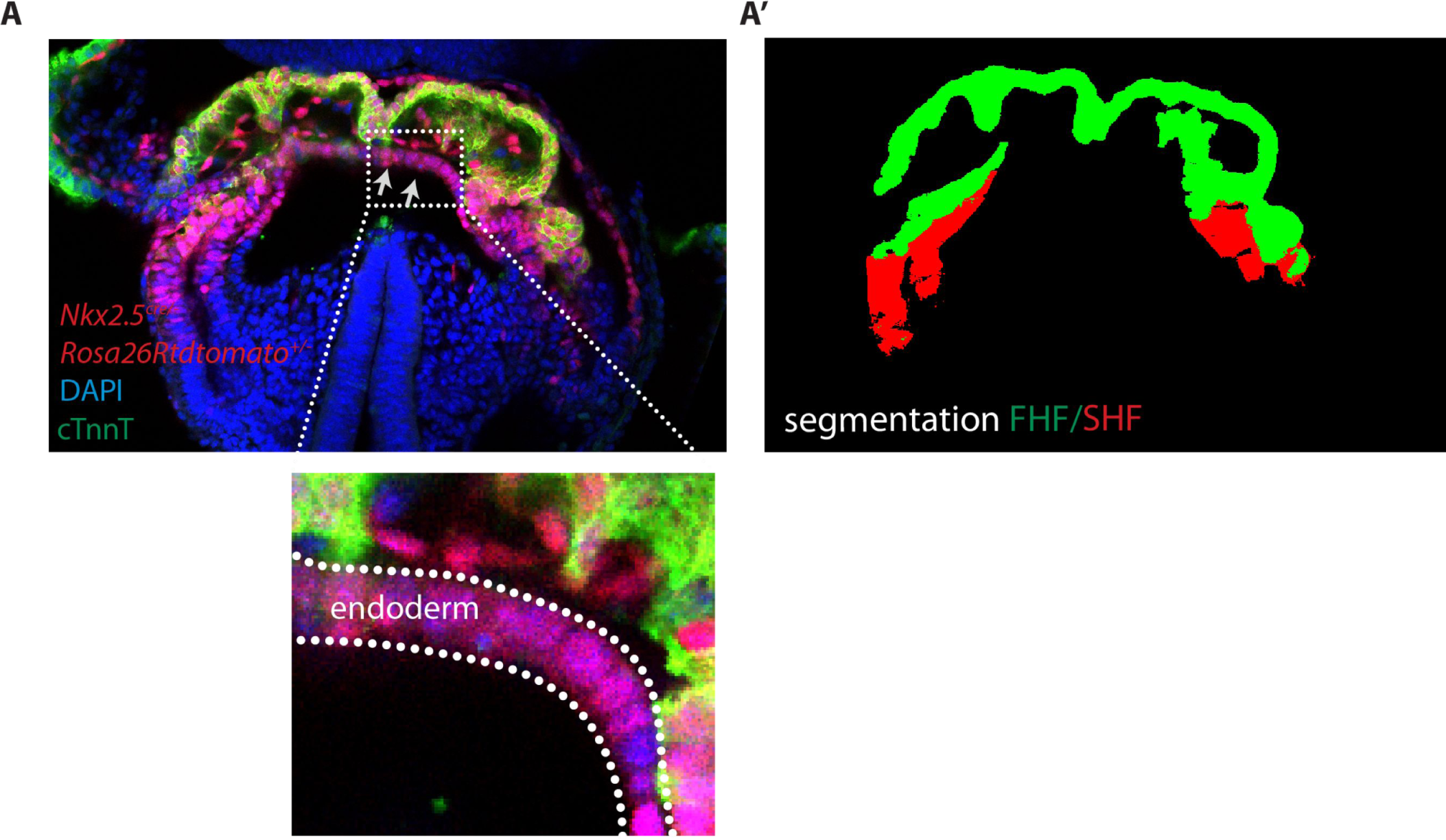
Image segmentation of the FHF and SHF. (A) Single optical section of an *Nkx2.5^cre/+^; Rosa26Rtdtomato^+/−^* embryo immunostained for cTnnT (green) and Dapi (blue). Gray arrows highlight tdtomato signal in the endoderm (A’) Segmentation of the FHF and SHF. Segmentation of the FHF (green) is based on the cTnnT signal while the segmentation of the SHF is based on the tdtomato positive and cTnnT negative signal (in red). tdtomato labeling in the endoderm and endocardium has been removed manually.

**Figure 2-figure supplement 2.**
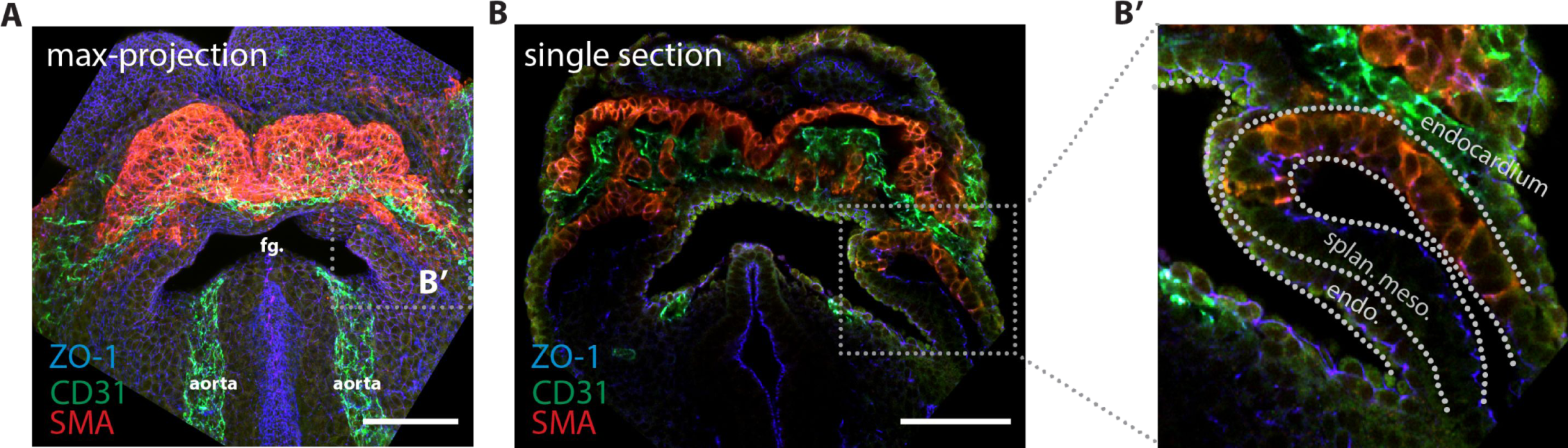
Endocardium localization in the cardiac crescent. (A) Maximum z-projection and (B, B’) single optical section of an embryo immunostained for smooth muscle actin (SMA, labeling differentiated cardiomyocytes) (red), CD31 (labeling the endocardium) (green) and ZO-1 (blue). endo: endoderm, splan. meso: splanchnic mesoderm Scale bars: 150μm. Figure Source Data 2 Source data for Figure 2C

**Figure 3-figure supplement 1.**
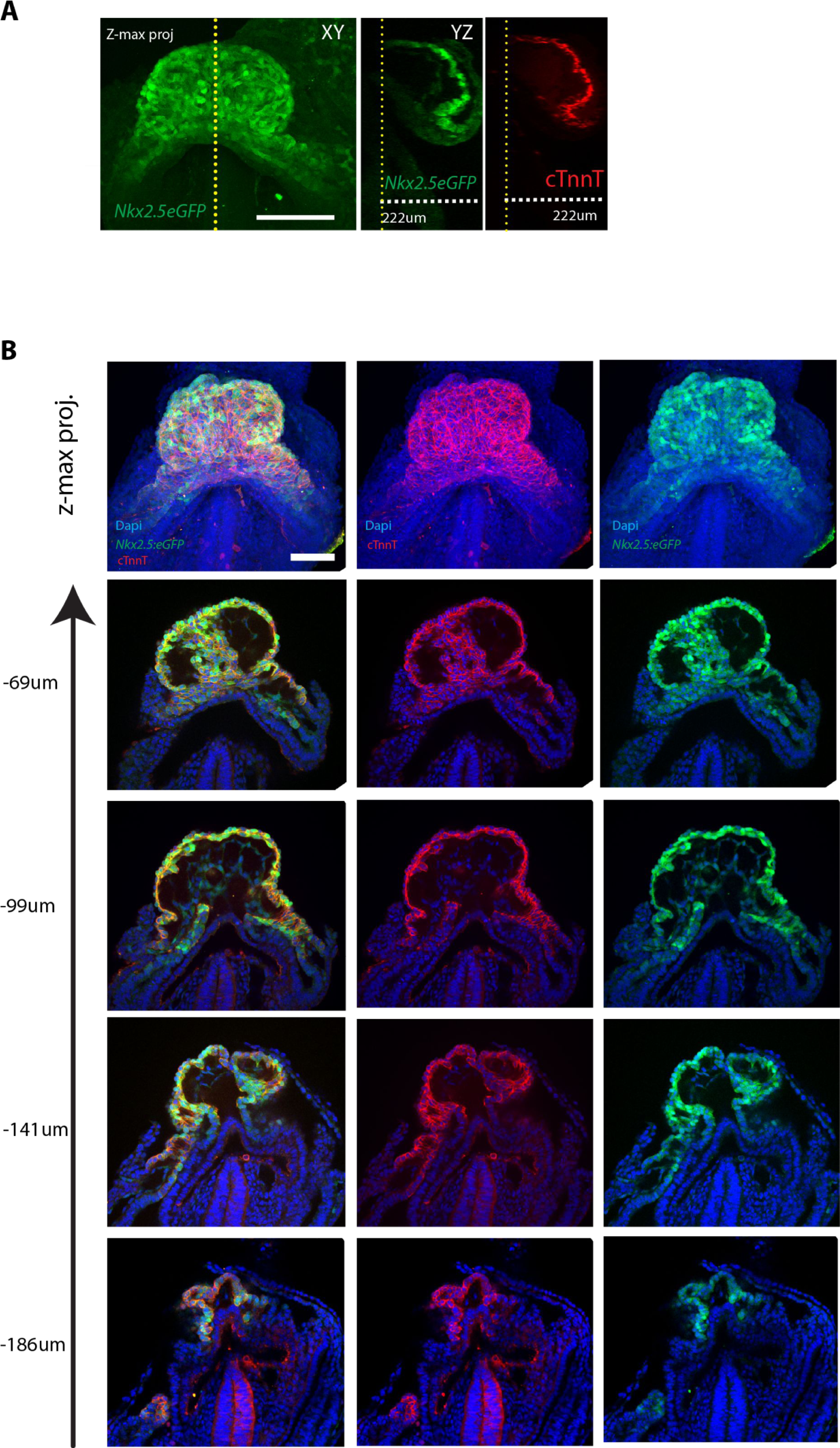
The *Nkx2.5eGFP* and cTnnT labeling at different z-levels. (A) z-maximum projection and yz view of a *Nkx2.5eGFP* embryo immunostained against cTnnT showing overlap between eGFP signal and cTnnT localisation along the z dimension. (B) z-maximum projection and optical sections at different z-level of an *Nkx2.5eGFP* embryo, immunostained for cTnnT (red) and Dapi (blue), showing overlapping expression of GFP and cTnnT signal. Scale bar: 100μm

**Figure 3-figure supplement 2.**
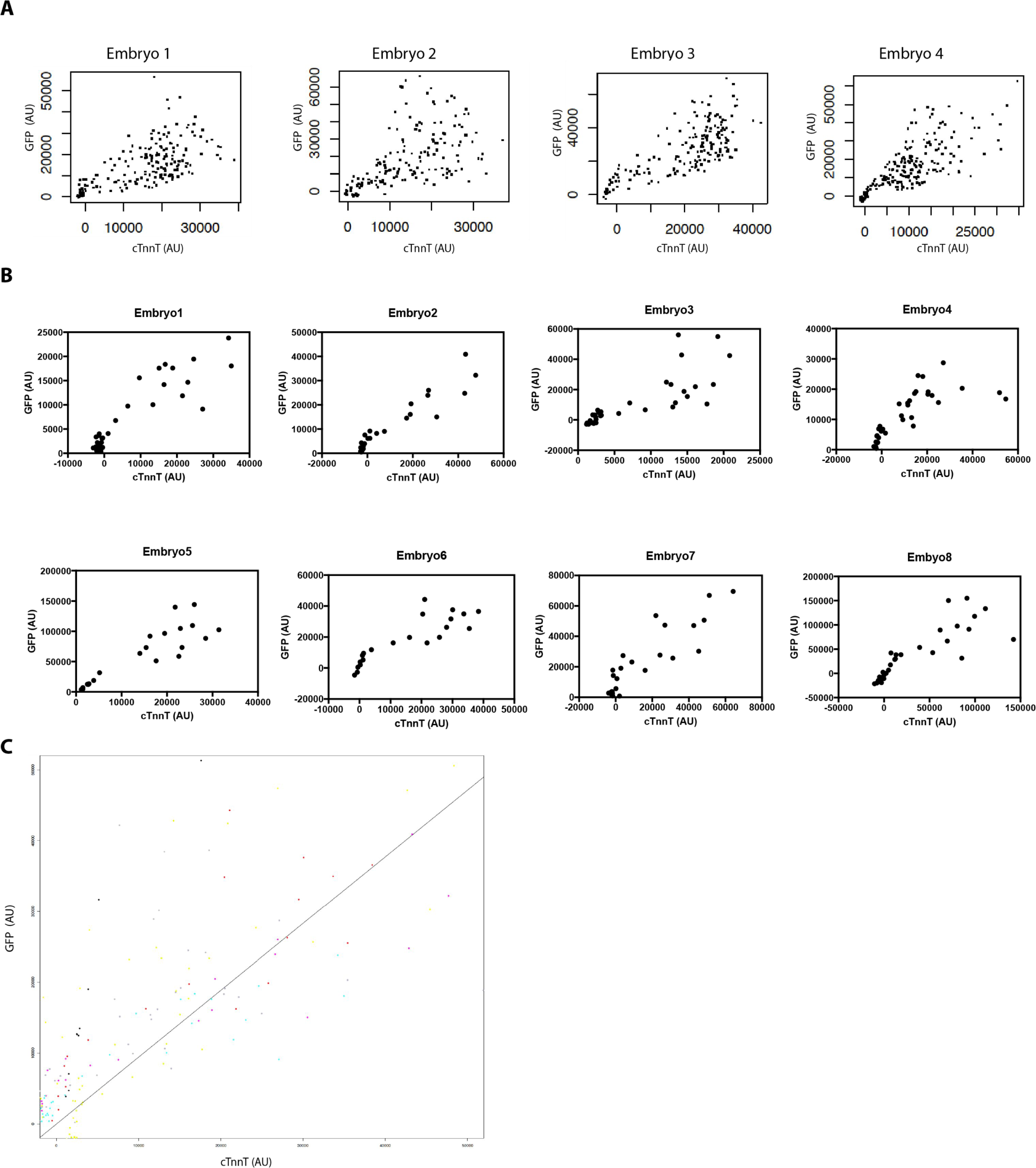
High GFP levels are detected in strongly labeled cTnnT cells. (A,B) Normalized GFP and cTnnT mean level for single segmented cell. Each dot represents a single segmented cell (A) located in the FHF and SHF and (B) located at the boundary zone between the FHF and SHF (n=130 cells analyzed from 8 embryos). (C) Linear mixed-effects model to find the relationship between the background substracted GFP and cTnnT levels adjusted by embryo for cell located at the boundary between the FHF and SHF (GFP=0.94*cTnnT, R2=0.77, p<2.2e-16). Cells are being considered positive for cTnnT when their mean intensity value is above 0. Note that small GFP signal can be detected in the cTnnT-negative SHF cells.

**Figure 3-figure supplement 3.**
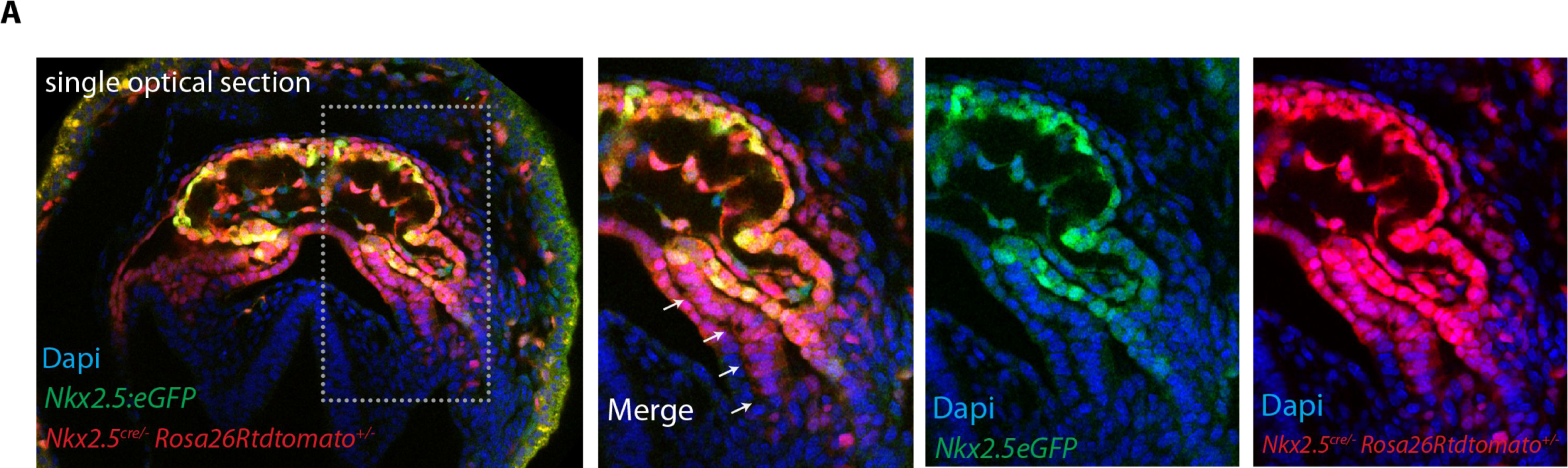
*Nkx2.5Cre* genetic tracing labels both the FHF and SHF. (**A**) The GFP expression of the *Nkx2.5eGFP* reporter does not fully overlap with tdtomato expression pattern in *Nkx2.5^cre/+^; Rosa26Rtdtomato^+/−^; Nkx2.5eGFP* embryos at the level of the splanchnic mesoderm/SHF. Scale bar: 100μm.

Figure Source Data 3

Source data for Figure 3D and E

**Figure 4-figure supplement 1.**
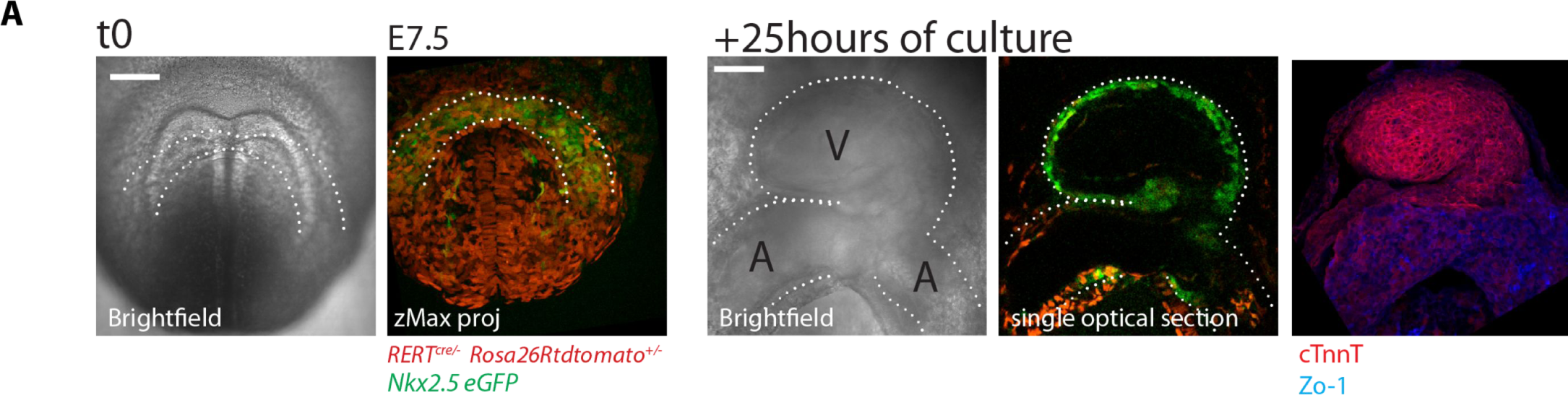
Embryos can be cultured and imaged under the 2-photon microscope for up to 24 hours. (A) *Nkx2.5eGFP*; *RERT*^+/−^; *Rosa26Rtdtomato*^+/−^embryo. Starting culture at E7.5, by the end of the 25 hours ex-vivo culture, the HT has formed and looped (experiment repeated 3 times independently). The embryo was subsequently fixed and immunostained for cTnnT (red) and ZO-1 (blue). V: prospective left ventricle, A: prospective atria. Scale bars: 100μm.

**Figure 4-figure supplement 2.**
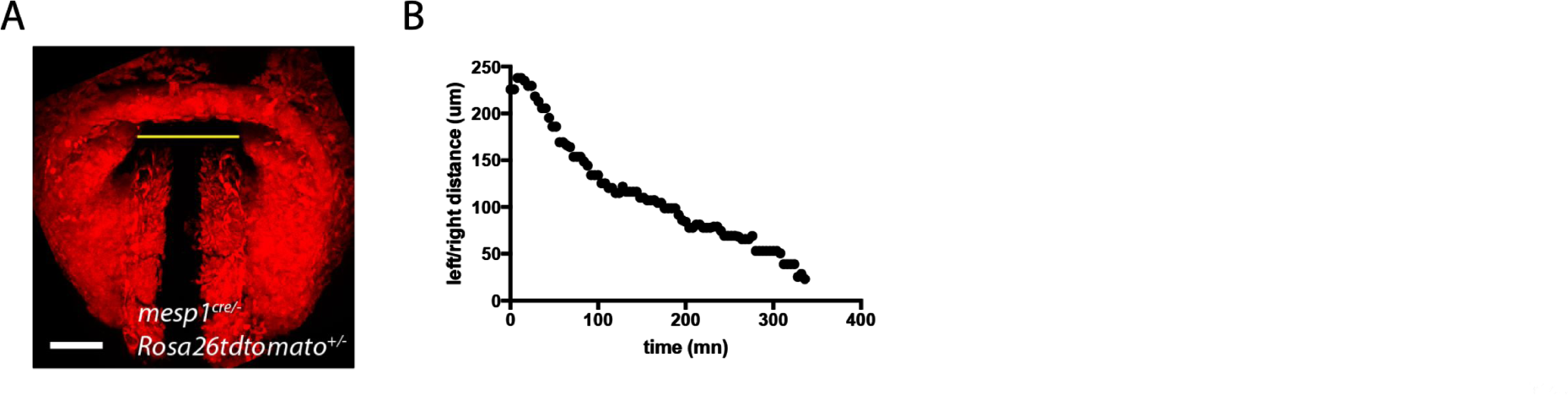
Quantification of the displacement of the splanchnic mesoderm during HT formation. (A) z-max projection of a *Mesp1^cre/+^; Rosa26Rtdtomato^+/−^* embryo at transversal HT stage. The yellow line indicates the distance between the left and right splanchnic mesoderm. (B) One example of the changes in left/right distance measured over time.

Figure Source Data 4

Source data for Figure Figure 4-figure supplement 2

**Figure 5-figure supplement 1.**
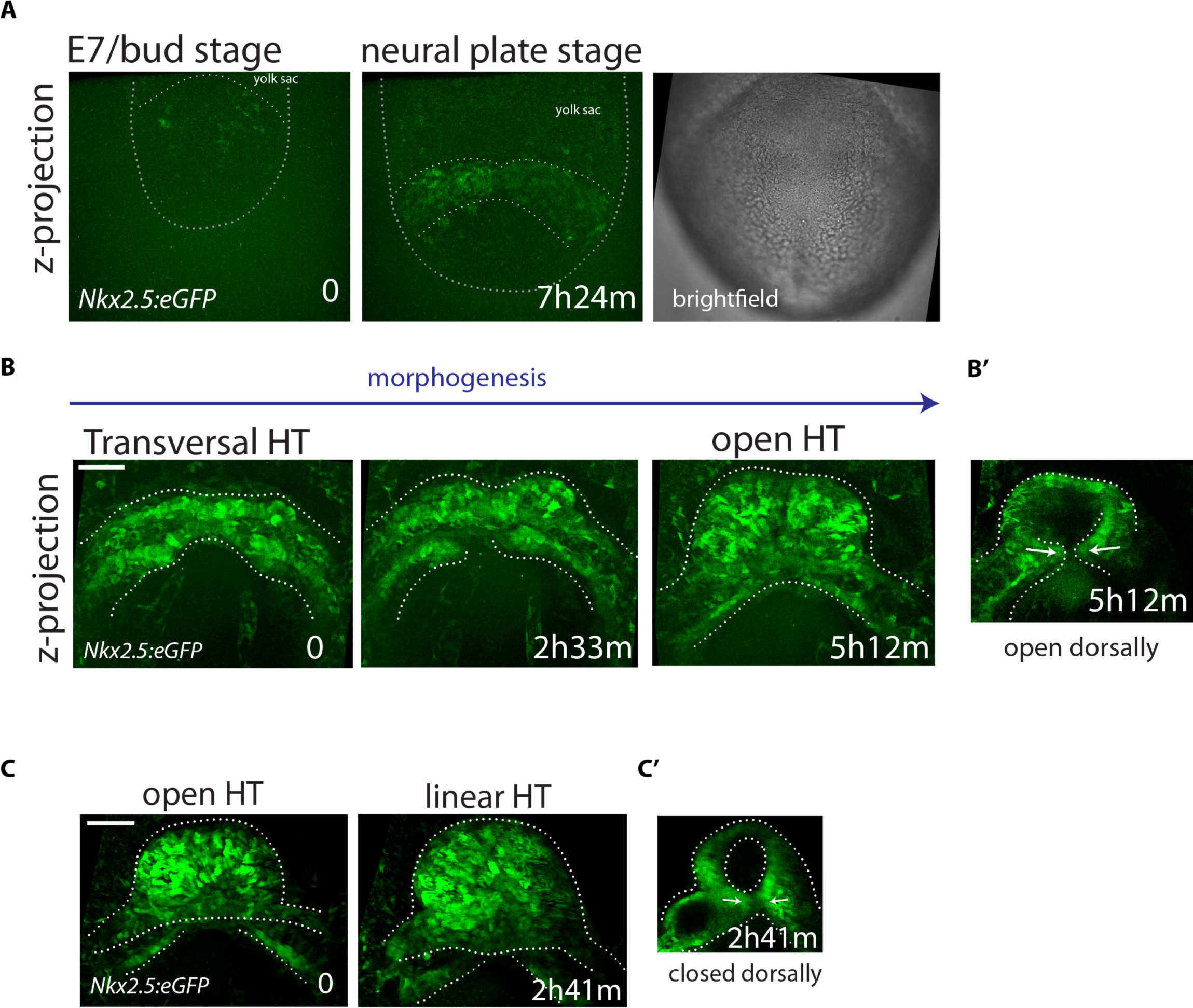
Live-imaging of *Nkx2.5eGFP* reporter line. (A-C) Time-lapse video sequences of *Nkx2.5eGFP* embryos from Early Bud (A), cc (B) and open HT stage (C). Images are z-maximum projections of 75 sections acquired every 4μm covering 300μm in (A), 55 sections acquired every 6μm covering 300μm in (B) and 59 sections acquired every 5μm covering 295μm in (C). (B’ and C’): z-max projection covering 132μm (B’) and 60μm (C’) showing the dorsal opening and the dorsal closure of the HT at the end of the videos (representative of 3 embryos). See also Videos 9 and 11. Scale bars: 100μm.

**Figure 5-figure supplement 2.**
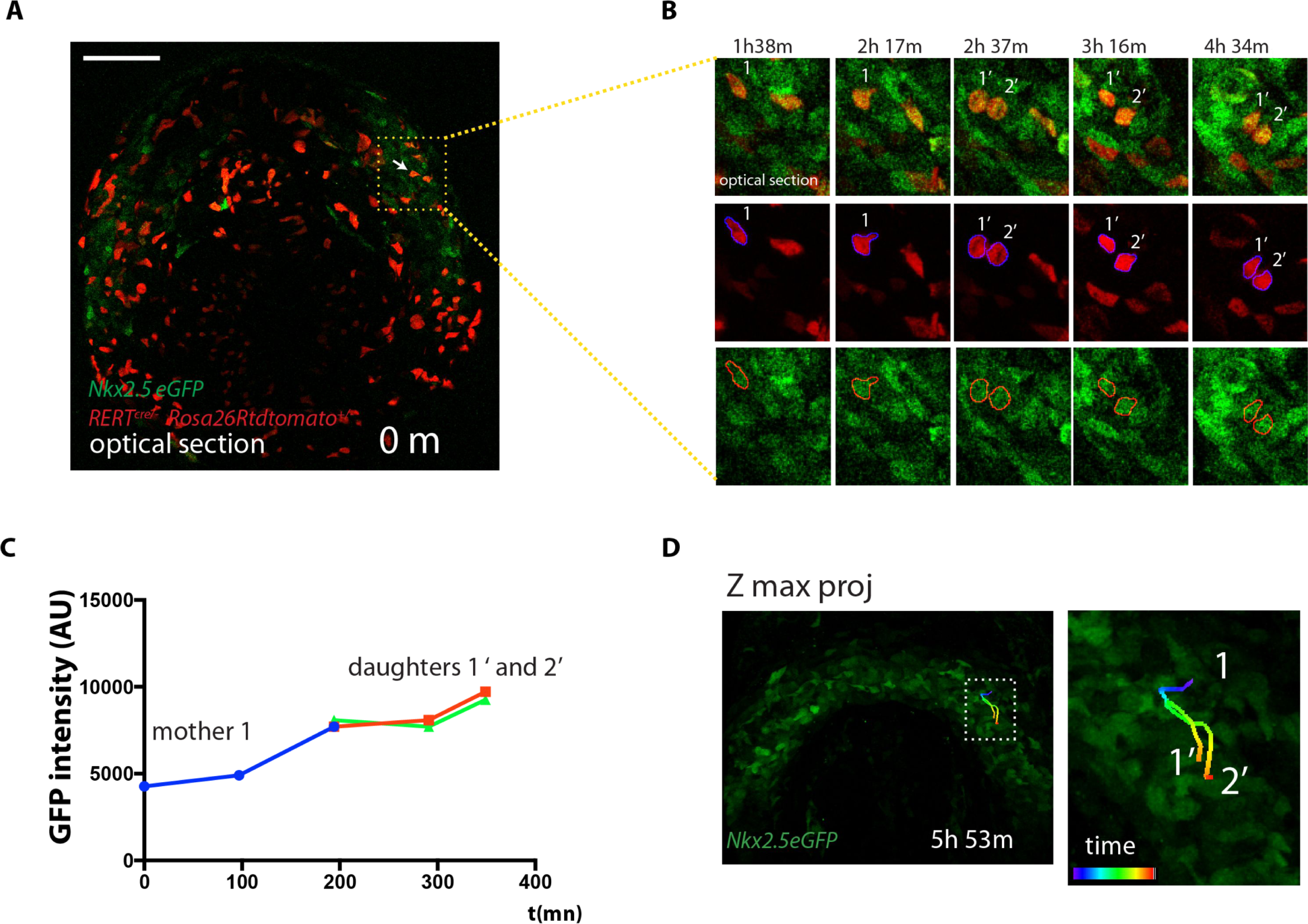
Cells divide during cardiac differentiation. (A-C) Time-lapse video of a dividing tdtomato+ cell. Images are single optical sections (same embryo as shown in Figure 4A). The dividing tdtomato+ cell and daughter cells are segmented and mean GFP level is measured. (D) Full 4D track of the dividing cell showing close localization of the progeny in the cardiac crescent. Scale bars: 100μm.

**Figure 5-figure supplement 3.**
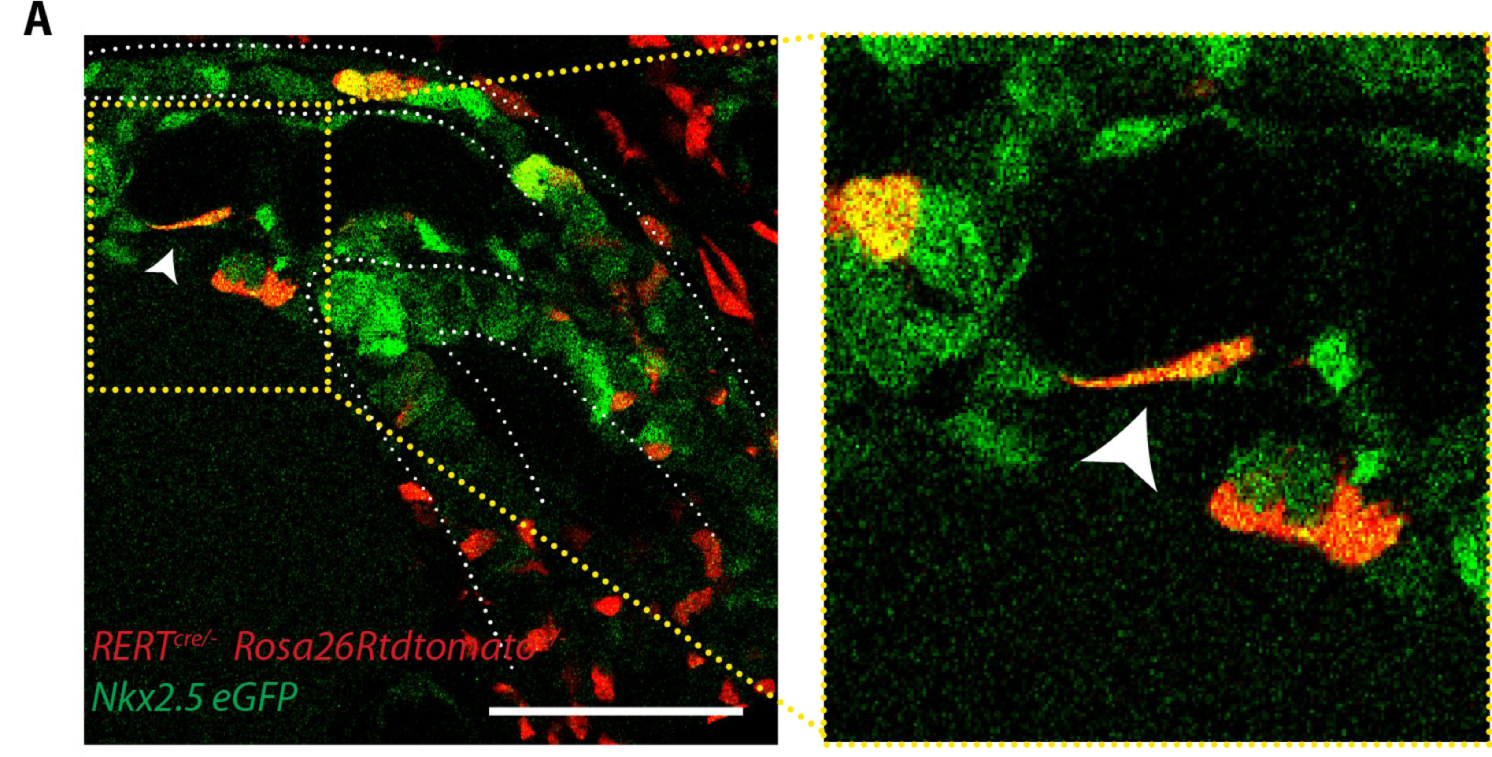
Endocardial cells show an elongated shape. (A) Single optical section of a tamoxifen-induced *Nkx2.5eGFP*; *RERT^+/−^; Rosa26Rtdtomato^+/−^* embryo showing a single endocardial labeled cell (white arrow). Scale bars: 100μm.

Figure Source Data 5

Source data for Figure 5A, D, F, G, I and Figure 5-figure supplement 1C, D

**Figure 6-figure supplement 1.**
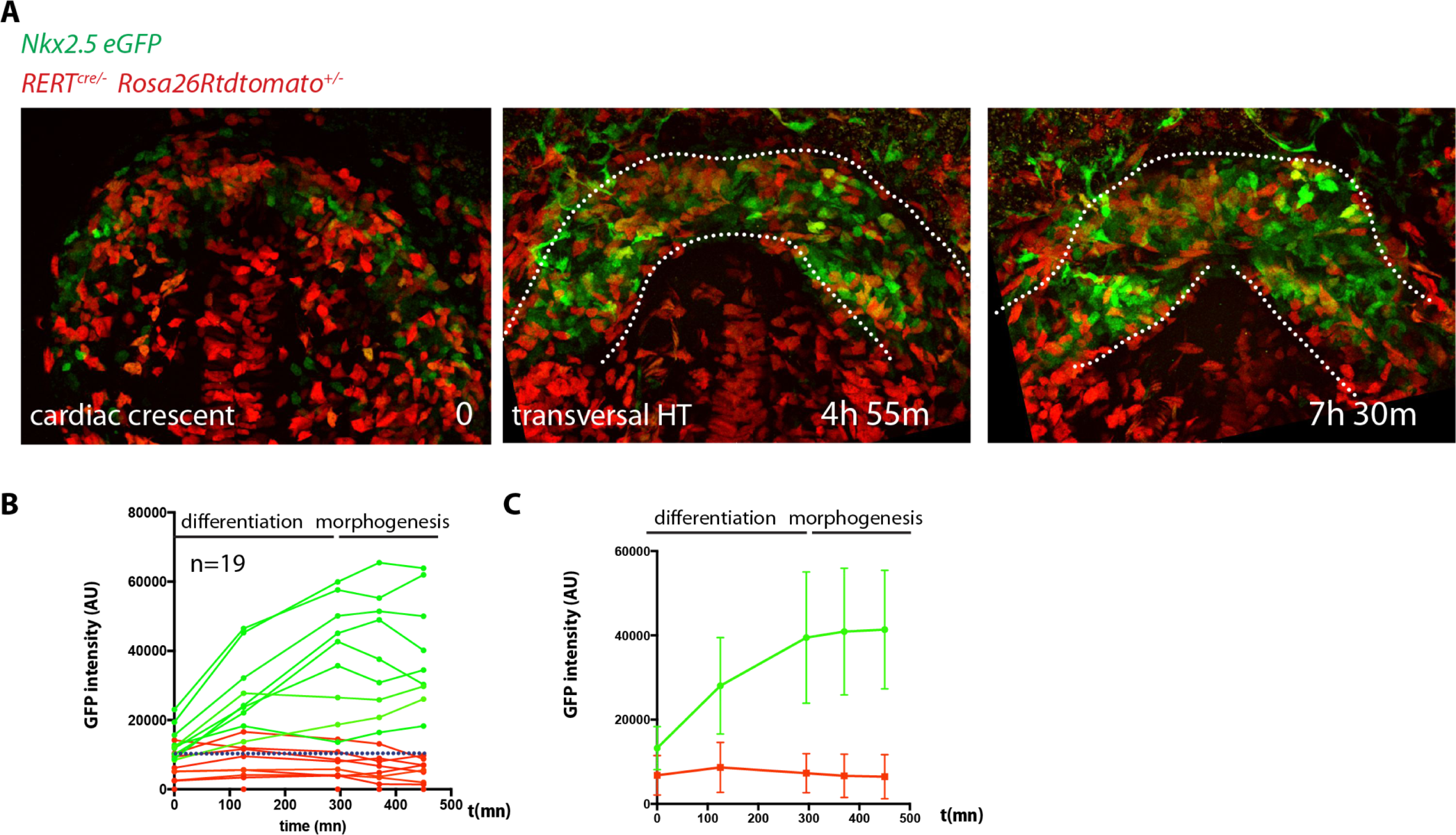
Longer videos can capture both initial cardiac differentiation in the FHF and early HT morphogenesis. (A) Initial and final images from a time-lapse video of a tamoxifen-induced *RERT^+/−^; Rosa26Rtdtomato^+/−^; Nkx2.5eGFP* embryo. Images are z-maximum projection of 44 sections acquired every 5μm covering 220μm. (B) GFP levels over time of the cells tracked in (A). GFP level for each tracked cell was measured at five successive time points. Green tracks represent cells with GFP intensity above the median intensity value of all the tracked cells at the last time point, while red tracks represent cells that maintain lower GFP levels. The blue dotted line represents the median intensity value. (C) Mean intensity value for all the green and red tracks and SD. Note that on average the GFP intensity of the green tracks becomes stable from around the transversal HT stage (around t=300m). Scale bars: 100μm.

Figure Source Data 6

Source data for Figure 6A, B, C, D, G, I, Figure 6-figure supplement 1A, Figure 6-figure supplement 1B and C.

**Figure 8-figure supplement 1.**
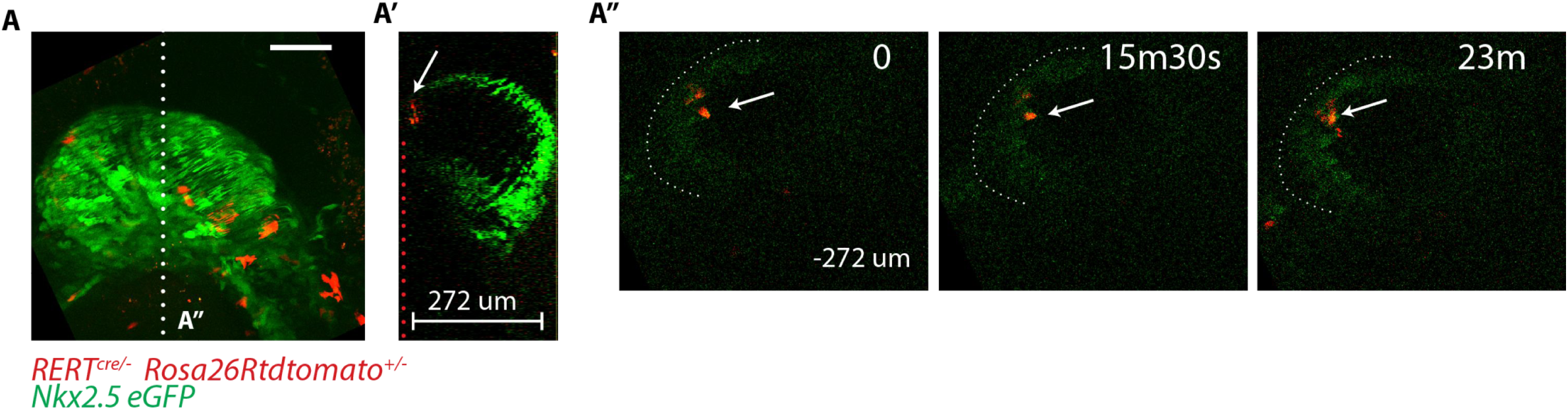
GFP level in deeper z levels cannot be accurately quantified. (A-A”) *RERT^+/−^; Rosa26Rtdtomato^+/−^; Nkx2.5eGFP* embryo at transversal HT stage. (A) z-maximum projection of 74 sections acquired every 4μm and covering 296μm. Same embryo as in Figure 6A. (A’) Single optical z-section positioned at the white dotted line shown in (A). This z-section shows the green signal decay at deep z positions (z= 0 to −274μm). (A”) An xy frame corresponding to a −272µm z-position (indicated by the red dotted line in (A’) at different time points during the video. The white arrow indicates td-tomato+ cells.

**Figure 8-figure supplement 2.**
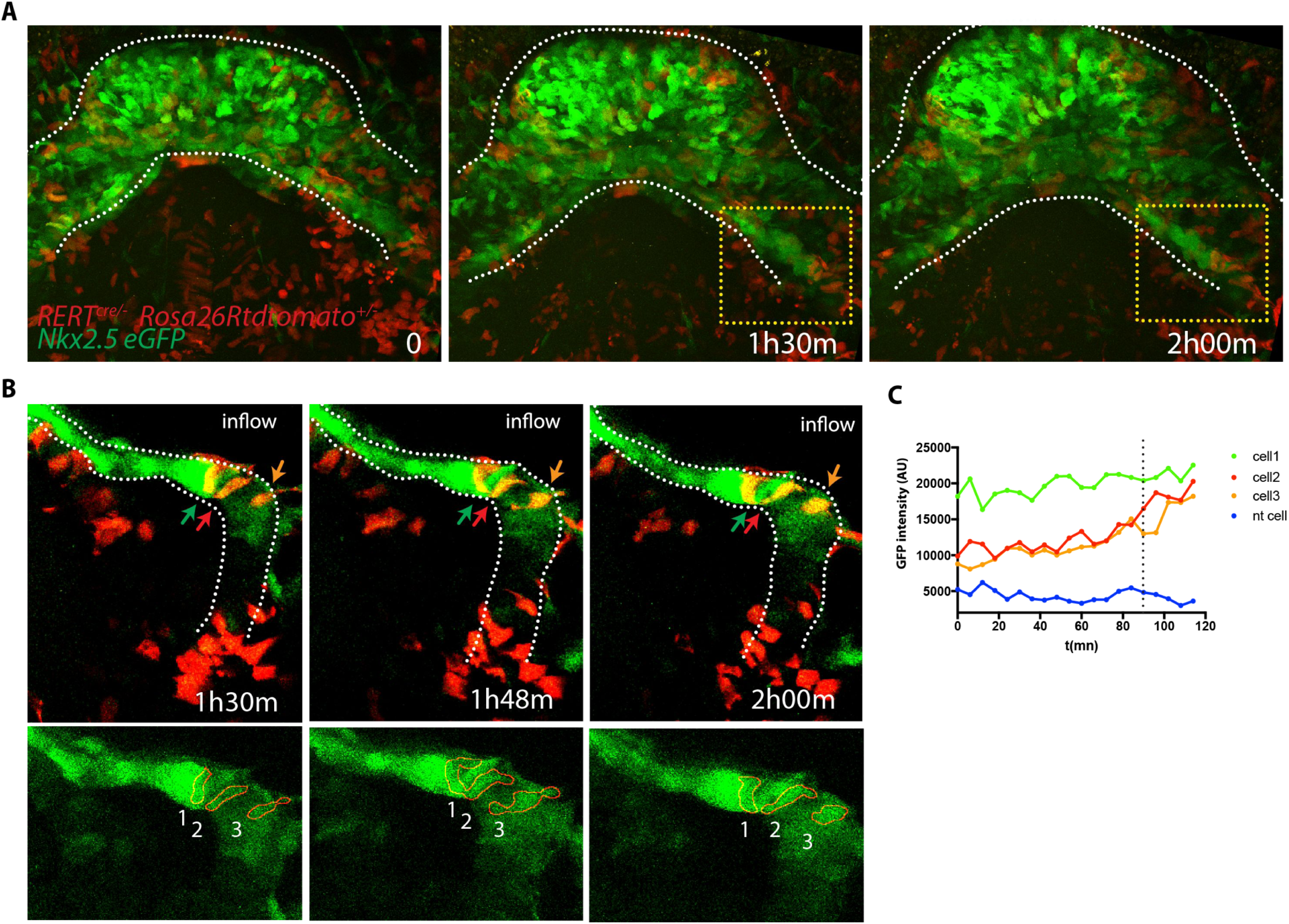
Cells increasing their GPF expression can be detected at the venous pole during activation of SHF contribution to the HT. (A) Time course of a tamoxifen-induced *RERT^+/−^; Rosa26Rtdtomato^+/−^; Nkx2.5eGFP* embryo during linear HT formation. Images are z-maximum projection of 44 sections acquired every 5μm covering 220μm. (B) Time course of individual cells tracked within the yellow-framed region in video (A). (C) GFP levels over time of the cells tracked in (B) Cell 1 shows high GFP level throughout the video while cell 2 and 3 increase their GFP level over time. The blue line represents a neural tube cell. The dotted line marks the 1h30m time point. Scale bars100μm.

Figure Source Data 8

Source data for Figure 8-figure supplement 2C.

## VIDEO LEGENDS

Video 1: 3D reconstruction of the cardiac crescent at EHF stage, based on *Nkx2.5^cre/+^ Rosa26Rtdtomato^+/−^* signal. Related to Figure 1A.

Video 2: 3D reconstruction of Mouse HT formation, based on embryos immunostained for cTnnT. Five representative stages are represented. Related to Figure 1C-G.

Video 3: 3D reconstruction of the FHF (green) and SHF (red). Based on *Nkx2.5^cre/+^ Rosa26Rtdtomato^+/−^* embryos immunostained for cTnnT. The border between cTnnT-negative and cTnnT-positive cells can be visualized at the interface between the red and green domains. Related to Figure 2B.

Video 4: z-maximum projection of an *Nkx2.5^cre/+^; Rosa26RmT/mG^+/−^* embryo at cc stage and single-section time-lapse video of the same embryo after 10h 42m of ex-vivo culture (representative analysis from 3 embryos). Interval between frames: 1s. Related to Figure 4E.

Video 5: Time-lapse video of a *Mesp1^cre/+^; Rosa26Rtdtomato^+/−^* embryo from cc up to open HT stage (h:mm:ss) (representative analysis from 4 embryos). Interval between frames: 4m 30. Duration of the video: 13h 16m 30s. Related to Figure 4F.

Video 6: Same embryo as in Video 5. Images are z-maximum projection of 9 sections acquired every 4μm covering 36μm (h:mm:ss) and allows visualization of the inside of the cardiac lumen during HT formation. Interval between frames: 4m 30s. Duration of the video: 8h 10m. Related to Figure 4F’.

Video 7: Time-lapse video of an *Nkx2.5^cre/+^; Rosa26RmT/mG^+/−^* embryo (h:mm:ss) (representative analysis from 2 embryos). Interval between frames: 22m. Duration of the video: 4h 02m. Related to Figure 4G-G’.

Video 8: Brightfield time-lapse video of a wild type embryo, from cc stage showing the onset of cardiomyocyte contractility and the formation of the cardiac lumen (h:mm:ss) (representative analysis from 2 embryos). Interval between frames: 1m 03s. Duration of the video: 6h 04m 21s. Representative analysis from 2 embryos.

Video 9: Same embryo shown in Video 14 zoomed-in at the level of the splanchnic mesoderm (mm:ss). SHF cells (green arrows) displace antero-medially relative to the underlying endoderm (red arrow). In addition, SHF cells move apart from each other (distance between the green arrows increase from 61 to 83μm over 3h12m). Related to Figure 6H and Figure 6-figure supplement 6A.

Video 10: Time-lapse video sequences of *Nkx2.5eGFP* embryos from Early Bud/E7.5 stage (h:mm:ss). (representative analysis from 2 embryos). Interval between frames: 12m. Duration of the video: 7h 24m Related to Figure 5-figure supplement 1A.

Video 11: Time-lapse video sequences of *Nkx2.5eGFP* embryos from cc to transversal HT stage (h:mm:ss). (representative analysis from 3 embryos). Interval between frames: 24m. Duration of the video: 6h 48m. Related to Figure 4H.

Video 12: Time-lapse video sequences of *Nkx2.5eGFP* embryos from transversal HT to open HT stage (h:mm:ss) (representative analysis from 3 embryos). Interval between frames: 8m 09s. Duration of the video: 5h 01m 33s. Related to Figure 5 - figure supplement 1B.

Video 13: Cell tracks in 3D+t during stages when cc differentiation takes place-from EHF onwards-(h:mm:ss). Cells are represented as green spheres if considered GFP+ at the end of the time-lapse analysis. and as red spheres when considered GFP-(Representative analysis from 3 embryos). Interval between frames: 19m 18s Duration of the video: 5h 28m 06s. Related to Figure 5A.

Video 14: Cell tracks in 3D+t during stages the open HT forms. Cells are represented as green spheres if considered GFP+ at the end of the time-lapse analysis and as red spheres when considered GFP-(Representative analysis from 3 embryos). Interval between frames: 7m 40s. Duration of the video: 3h 12m. Related to Figure 6A.

Video 15: Time-lapse video of a *RERT^+/−^; Rosa26Rtdtomato^+/−^; Nkx2.5eGFP* embryo during the stages at which the transversal HT transforms into the open HT (h:mm:ss). Interval between frames: 19m 19s. Duration of the video: 7h 24m 35s. (2nd half) 3D reconstruction at three time points based of the eGFP signal (green) and red-labeled cells located in the splanchnic mesoderm. Related to Figure 7A.

Video 16: Time-lapse video of an *Isl1^cre/+^; Rosa26Rtdtomato^+/−^; Nkx2.5eGFP* embryo (h:mm:ss) (representative analysis from 3 embryos). Interval between frames: 6m. Duration of the video: 7h 18m. Related to Figure 8A.

Video 17: Same embryo shown in Video 16 zoomed-in at the level of the splanchnic mesoderm (h:mm:ss). Related to Figure 8C.

Video 18: 3D rendering of an *Isl1^cre/+^;Rosa26Rtdtomato^+/−^; Nkx2.5eGFP* embryo at transversal HT stage. Related to Figure 9A.

Video 19: 3D rendering of an *Isl1^cre/+^;Rosa26Rtdtomato^+/−^; Nkx2.5eGFP* embryo at linear HT stage. Related to Figure 9B.

